# The Chromosome 19 miRNA Cluster Guards Trophoblasts Against Overacting Innate Immunity

**DOI:** 10.1101/2025.04.03.647038

**Authors:** Jean-Francois Mouillet, Yingshi Ouyang, Elena Sadovsky, Vinoth K. Kothnadan, Heather L. Sorenson, Logan J. Badeau, Saumendra N. Sarkar, Tianjiao Chu, Alexander Sorkin, Yoel Sadovsky

## Abstract

To maintain pregnancy health, the human placenta delicately balances protection of the developing fetus from invading pathogens with suppression of excessive inflammation that could lead to fetal and neonatal autoimmune disorders. Previous research, including our own, has shown that small RNA products of the Chromosome 19 MicroRNA Cluster (C19MC) promote viral resistance in non-trophoblastic cells. However, the role of C19MC products in placental trophoblasts remained unclear. Here, we analyzed chromatin accessibility in the C19MC enhancer and identified a previously unknown regulatory domain. Deletion of this domain silenced the expression of C19MC microRNA and Alu elements in trophoblasts. This silencing unexpectedly led to marked activation of cellular innate immune response and strikingly increased Toll-like receptor 3 (TLR3)-mediated sensitivity to poly(I:C), a viral RNA mimic. Our data suggest that C19MC non-coding RNAs interfere with endosomal TLR3 activation in trophoblasts, highlighting a previously unrecognized mechanism for hindrance of excessive innate immune activation.

## Introduction

The placenta plays an indispensable role in supporting Eutherian pregnancy. From implantation and throughout pregnancy, the placenta constitutes the maternal-fetal interface, where it promotes fetal development and growth by regulating the transfer of gases and nutrients to the developing embryo and the removal of waste products^1^. Furthermore, the placenta produces hormones, chemicals, growth factors, and extracellular vesicles (EVs) that mediate maternal- fetal communication and adaptation to pregnancy, and provides a balanced immune defense against pathogens, thus safeguarding the health of the mother and the future offspring ^2,3^.

Consequently, placental dysfunction is a major contributor to the pathology of common pregnancy complications, including fetal growth restriction, preeclampsia, pregnancy loss, and related disorders^4^.

Within the human placenta, embryo-derived trophoblasts form the multinucleated interface that directly contacts maternal blood and uterine decidua. This interface governs the placenta’s critical transport, metabolic, and immune functions^5^. Human trophoblasts, from as early as the first trimester of pregnancy, have been defined^6^ by (a) the expression of cytokeratin 7, a key trophoblast epithelial marker^7^; (b) hypomethylation of the Elf5 promoter, which leads to enhanced Elf5 expression and the promotion of trophoblast development^8^; (c) absent expression of major HLA Class I molecules HLA-A, and -B, which reduces placental immunogenicity of the semi-allographic fetus^9^; and (d) the expression of primate-specific microRNAs (miRNAs) of the Chromosome 19 miRNA cluster (C19MC)^10^. The role of C19MC RNAs has been puzzling. Largely placenta-specific and vastly abundant in human trophoblasts, the C19MC’s 59 miRNA species are expressed from 46 miRNA genes that are scattered over a region of nearly 100 kb. These miRNAs are intermingled with numerous Alu-type short interspersed nuclear elements that exhibit a preferred antisense orientation^10–14^. C19MC miRNA are released to the maternal circulation within the first 2 weeks of implantation, when their level rapidly rises and then remains stable across human pregnancy^15^.

Several functions for C19MC ncRNAs have been posited. The rare extra-placental expression of C19MC miRNA in primitive neuroectodermal brain tumors has been associated with aggressive tumor behavior, with several C19MC members identified as oncogenes^16^ (and reviewed in ^17^). Several other tumors have been associated with C19MC expression ^18,19^. We previously showed that the expression of C19MC in non-trophoblastic cells or the exposure of non-trophoblastic cells to C19MC-containing trophoblastic small EVs augmented the resistance of these cells to viral infections, an effect instigated by enhanced autophagy^20^. Further research revealed that C19MC Alu-RNAs trigger innate immunity in non-trophoblastic cells by stimulating the RIG-I pathway^21^, leading to the production of interferons (IFNs), particularly IFN lambda (IFNL), which is typical of the placental interferon response^22^. While previous studies examined C19MC RNAs in other cell types, their function in placental trophoblasts, where they originate, was unknown. We comprehensively analyze the C19MC’s regulatory mechanisms and role in trophoblasts, and identified a specific enhancer element. Deletion of this element led to complete silencing of the C19MC region and an unexpected, robust stimulation of trophoblast innate immunity through the TLR3 pathway. This suggests that C19MC normally acts to guard trophoblasts against excessive innate immune activation.

## Results

### Identification of ATAC-positive C19MC open chromatin domains that exhibit autonomous enhancer activity in vitro and in vivo

We previously showed that a BAC plasmid harboring 160 kb of the human C19MC genomic locus was sufficient for effective heterologous expression of the C19MC miRNA^23,24^. The BAC contained 60 kb sequences upstream of the first miRNA gene, which included a CpG island, previously shown to exhibit promoter activity^25^, and excluded the small miR-371 cluster, located downstream. These data implied that the regulatory domains that are essential for C19MC expression were present within the BAC sequences. To identify open C19MC chromatin regions within the C19MC locus and examine their tissue-specific activity, we deployed ATAC-seq^26^, and identified 19 trophoblast-specific, enriched accessible chromatin peaks in placenta-derived primary human trophoblasts (PHT cells) that were absent in C19MC non-expressing HUVEC cells (Fig.1a). Reassuringly, peak 9 (ATAC-9) aligned almost perfectly with the previously identified CpG^25^, yet our comprehensive approach suggested the presence of additional C19MC regulatory elements.

To determine if ATAC-seq peak regions harbor autonomous enhancer activity, we cloned each of the 19 ATAC-positive peak sequences upstream of the SV40 minimal promoter in a pGL3 vector. Due to the proximity of ATAC-3 and -4 peaks, as well as ATAC-13 and -14 peaks, we cloned two segments that each spanned two adjacent peaks (denoted ATAC-3/4 and ATAC- 13/14, respectively), and transfected each reporter into PHT cells and non-placental cells. As shown in Fig. 1b, four of the 19 constructs consistently exhibited increased luciferase activity across *all* C19MC-expressing trophoblast cells (BeWo, JEG3, and PHT). However, they were inactive in non-placental 293T and HeLa cells and in the extravillous trophoblast-derived cell line HTR-8/SVneo, which expresses very low levels of C19MC^24,27^. Because each construct contained the ATAC-seq regions along with longer flanking sequences to account for imprecise localization of the predicted peak, we cloned a shorter version of the four regions (ATAC-peak 3/4, ATAC-11, ATAC-12, and ATAC-17), and validated their luciferase activity, including their activity when cloned in reverse orientation, as characteristic of enhancer regions (Fig. S1a-b). These shorter constructs were used in subsequent experiments. Note that ATAC-9, which harbored the CpG island, was not sufficient to enhance transcriptional activity in our heterologous reporter assay, suggesting that this ATAC9 element may function as a promoter rather than an enhancer, consistent with the behavior of some regulatory elements^28^.

**Fig. 1:**
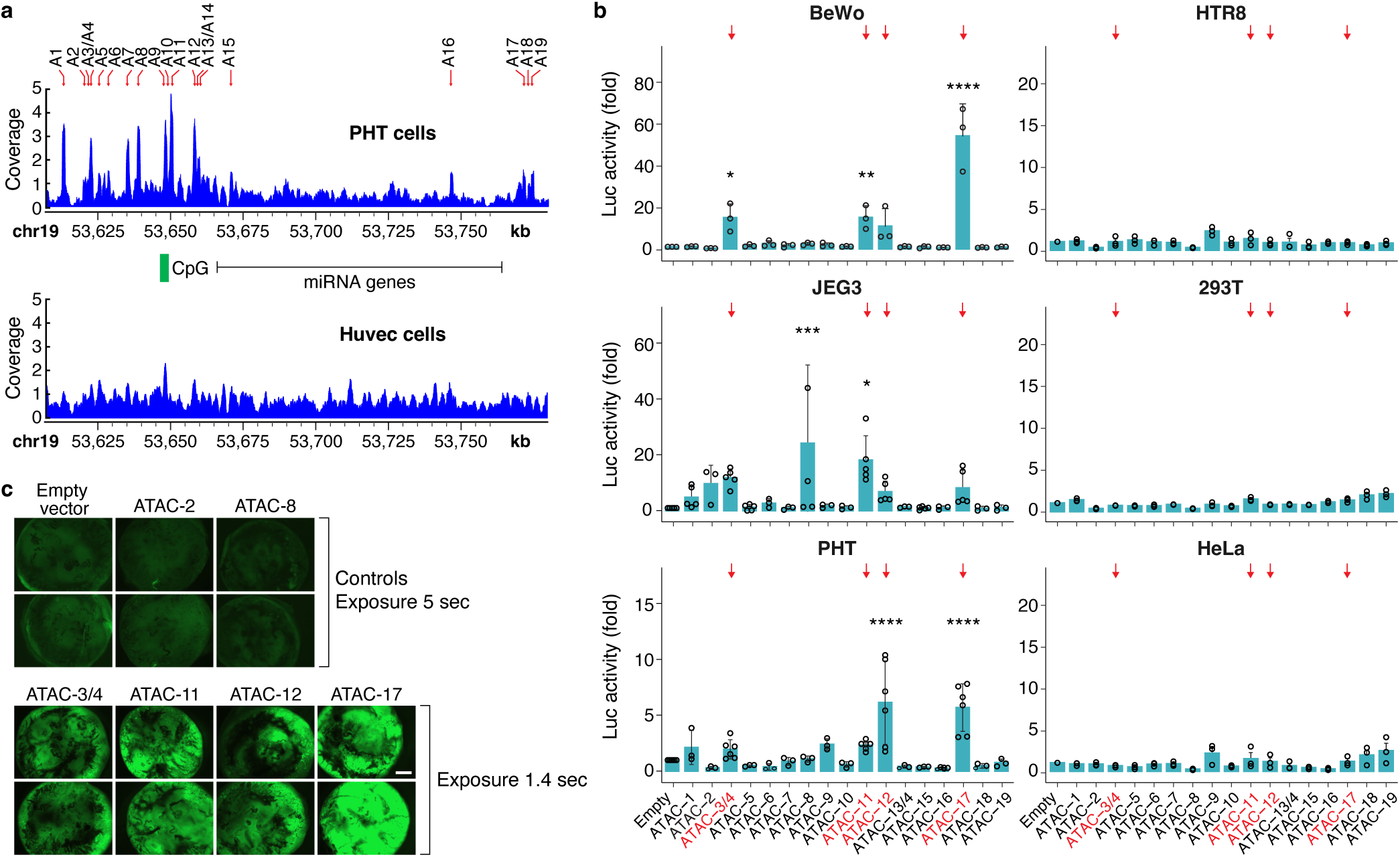
Identification of open chromatin, transcriptionally active sites in the C19MC region. **a.** Sequencing track for the C19MC locus, showing the ATAC-seq peaks in primary human trophoblasts (PHT) and in HUVEC cells. The red arrows indicate the location of statistically different peaks between PHT and HUVEC cells. The green box indicates the location of the previously identified CpG island. **b**. Functional enhancer assays using a luciferase reporter. Sequences corresponding to the ATAC peaks were cloned in the pGL3- reporter plasmid and transiently transfected along with a *Renilla* luciferase plasmid, used for normalization. Note that the relatively low activity detected in PHT cells is due to the very low transfection efficiency in primary trophoblasts. The orange arrows indicate the enhancer regions that exhibit transcriptional activity in all C19MC-expressing trophoblasts and that were selected for further investigation. Data are mean ±SD (n=3-6). * p<0.05, ** p<0.01, *** p<0.005, **** p<0.0001. **c**. The activity of ATAC-positive regions *in vivo*. ATAC sequences were cloned into a pLV-hsp68-eGFP lentiviral vector^29^. Lentiviruses were produced and used to transduce blastocyst stage embryos for 5 h before transplanting them into pseudo-pregnant females. The successful integration of the transgene was verified by PCR on placental genomic DNA. Placentas were recovered 14 days after transfer and fluorescently imaged. Placentas expressing negative control constructs (upper panel) were exposed for a longer time to visualize the placentas. Scale bar =1 mm.

To validate our results *in vivo*, we tested the activity of the four identified ATAC-peak elements selectively in the mouse placenta, using trophoblast-specific lentiviral gene transfer, as we previously showed^29^. We cloned individual ATAC elements into the transgenic reporter plasmid hsp68-eGFP (a gift to Addgene from Benjamin Yu, Dermatology, UC San Diego), and found that the four regions, identified on the basis of our *in vitro* studies, robustly drove GFP expression in the mouse placenta (Fig. 1c). Together, these experiments confirmed the presence of trophoblast-specific autonomous enhancer activity in these four C19MC regions.

### The function of ATAC-positive domains in the context of intact chromatin

We next sought to corroborate the enhancer activity of these four enhancer regions in the context of their native chromatin environment. We chose to edit BeWo cells, because (a) these cells exhibit the characteristic differentiation and fusion features of placental trophoblasts^6,30^, (b) they express high levels of C19MC miRNAs^31^, and (c) unlike PHT cells, BeWo cells are accessible to CRISPR-*Cas9* editing. Using a pair of guide RNAs flanking each targeted region, we used *Cas9* to delete both enhancer alleles for these elements. Remarkably, using two independent ATAC-11 deletion clones (herein referred to as BeWo-dA11.21 and -dA11.23 clones), we found that deletion of this region, but not those lacking the other ATAC elements, led to a very low C19MC ncRNA output (Fig. 2a and Fig. S1c). Although we could not generate a homozygous deletion of the ATAC-17 enhancer, deletion of one allele did not impact C19MC expression. In ATAC-11 deletion clones, the expression of a non-C19MC member, miR-21, was higher, possibly due to reduced concentration of C19MC substrate for the miRNA processing enzymes. RNA-seq analysis of the two ATAC-11 deletion clones, with an emphasis on a 1.5 Mb region around the C19MC locus, revealed a marked reduction in Alu and non-Alu RNA reads from the C19MC region (Fig. 2b-c), with no significant expression changes outside the C19MC region (Fig. 2d), supporting the specific role of the ATAC-11 domain in regulation of the C19MC’s ncRNA output.

**Fig. 2:**
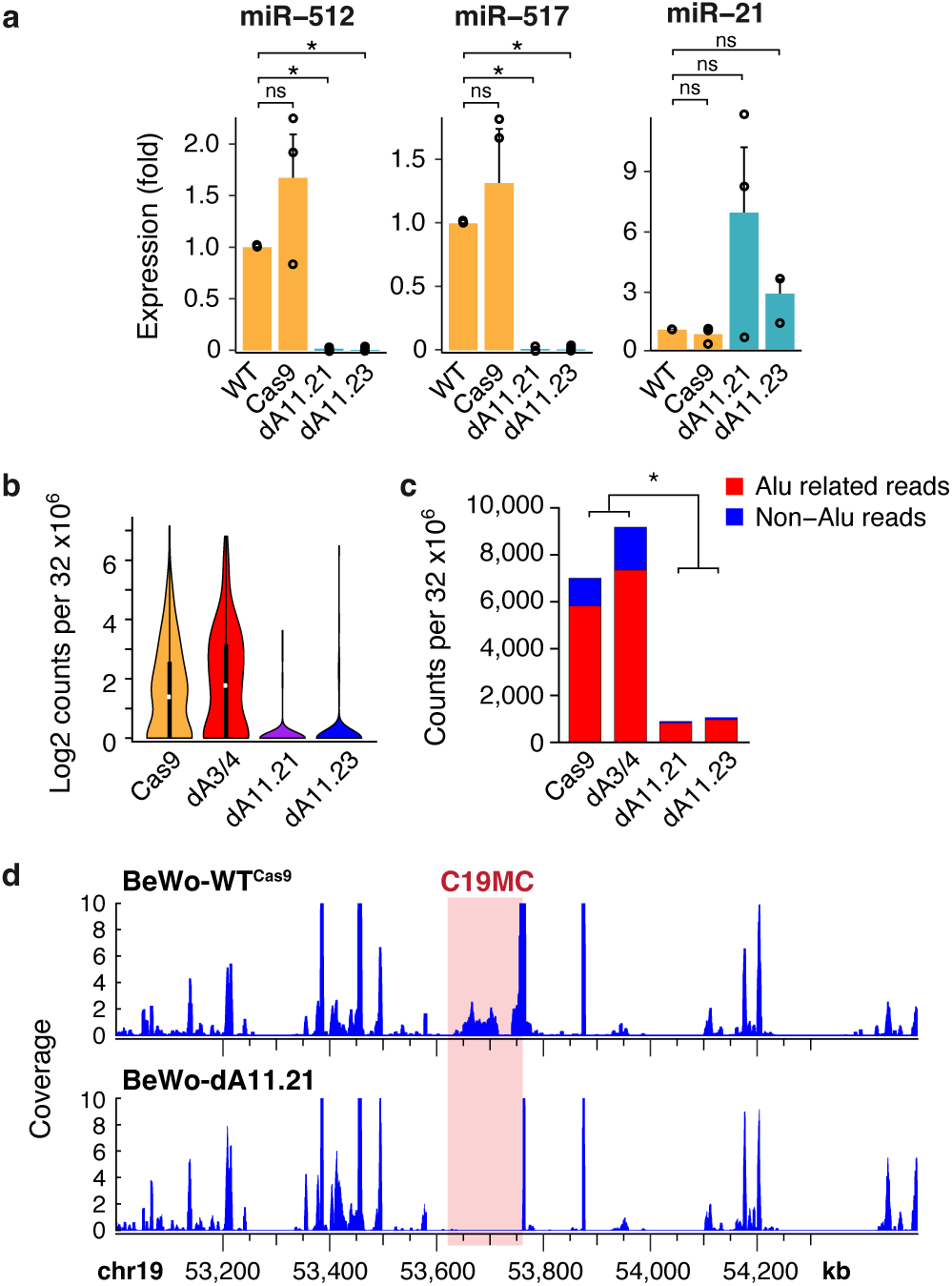
C19MC small RNA output in the C19MC-dA11-deleted cells. **a.** RT-qPCR analysis of miRNA expression in BeWo cells, showing a sharp reduction of C19MC miRNA expression (miR-512 and miR-517) and increased expression of a non-C19MC miRNA (miR-21). Four lines were tested, including BeWo-WT, BeWo cells transduced with a doxycycline-inducible Cas9 lentivirus (Cas9), and two independent clones with a homozygous deletion of ATAC-11 (BeWo-dA11-21 and -dA11-23). Data are mean ±SD, n=3. * p<0.05; ns = nonsignificant. **b**. The expression of C19MC Alu RNAs in BeWo-Cas9 or -dA11 (2 lines each), determined by RNA-seq (4 libraries). In the left panel, reads are Log2 expression of Alu elements, normalized to library size. The line dA3/4 was Cas9-edited at ATAC-3/4 site, as an additional control. Y axis: Log2 expression of Alu elements, normalized by library size. The expression difference between each control group and each deletion group was significant at p < 2.2x10^-16^. **c.** The normalized total counts for all Alu RNAs (red) and all non-Alu small RNAs (blue, including miRNA) in the same samples as in b. The difference between the fraction of total Alu and non-Alu RNAs in Cas9 and dA3/4 vs the two dA11 clones was significant (p<0.05, t-test on log fraction). **d**. An RNA-seq density plot, representing the chromosome 19 region that harbors the C19MC locus, plotted along chr19:53000000−54500000. Each bar represents the log2 read frequency, plotted by the chromosome coordinates. The pink box indicates the C19MC location.

To ensure the absence of major genomic disruption in the edited BeWo clones, we performed a cytogenetic and FISH analyses of the ATAC-11 clones *vs* BeWo-WT and found no significant differences (Fig. S2a). In particular, we observed three copies of chromosome 19 and its C19MC locus in both control and ATAC-11-edited BeWo cells, known to have a triploid genome. Moreover, the ATAC-11-edited BeWo cells exhibited similar chromosome counts and no significant anomalies upon genomic analysis assay of control BeWo-Cas9 and -dA11 cells (Fig. S2b-c).

### Amplified innate immunity and viral resistance in C19MC-depleted cells

In light of our previous observations^20^, we tested our C19MC-depleted cells for their resistance to infection by VSV. Unexpectedly, we found that BeWo-dA11 were more resistant to infection than the parental BeWo-WT cells (Fig. 3a). Correspondingly, using RNA-seq, we found marked upregulation of IFNs and IFN-stimulated gene (ISG) expression in BeWo-dA11 (Fig. 3b), which we confirmed using enrichment analysis for biological processes with the clusterProfiler R-package or Ingenuity Pathway Analysis (IPA, Fig. S3a-b). To induce an innate immune response similar to a viral infection, we exposed both control and dA11 cells to poly(I:C), a synthetic double-stranded RNA analog^32^. As shown in the volcano plots in Fig. 3c, we found that BeWo-dA11 cells exhibited marked upregulation of ISGs when compared to WT cells, and this response was strikingly amplified in the presence of poly(I:C).

**Fig. 3:**
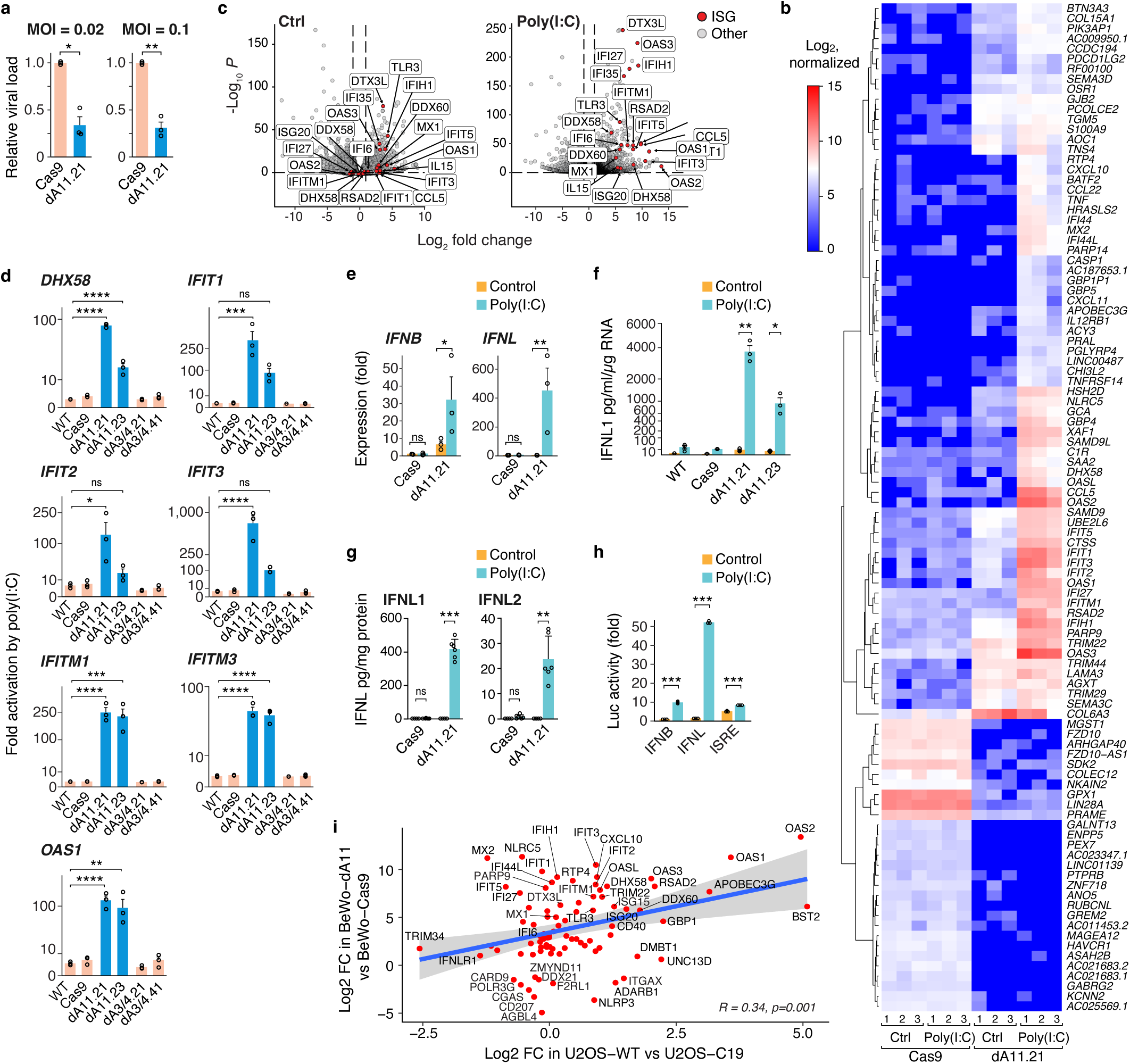
C19MC-silenced cells and innate immune response. **a.** Viral resistance in BeWo- dA11 cells. The cells were inoculated with the VSV virus at multiplicity of infection (MOI) of 0.02 and 0.1 for 5 h. The viral load was assessed by RT-qPCR of the viral protein VSV-G. Data shown as mean ±SD. * p<0.02; ** p<0.001, n=3. **b**. A heat map of the 100 top differentially expressed genes in BeWo-Cas9 *vs* -dA11, unstimulated (Ctrl) or exposed to 10 μg/ml poly(I:C) overnight. Three libraries per line were sequenced. **c**. Volcano plots depicting differentially expressed genes in BeWo-Cas9 *vs* -dA11, unstimulated (left panel) or exposed to 10 μg/ml poly(I:C) (right panel). **d**. RT-qPCR analysis for selected ISGs. The bars represent the fold induction by poly(I:C) (10 µg/ml). Data are displayed using a square root scale y-axis to account for the large differences between the cell lines. Data are mean ±SD, n=3. * p<0.05, ** p <0.001, *** p<0.0005, **** p<0.0001, ns = not significant. **e**. RT-qPCR analysis for interferon-β (IFNB) and interferon-λ (IFNL). Cells were stimulated by 10 µg/ml of poly(I:C). Data are mean ±SD, n=3, *p<0.05, ** p <0.001. **f**. IFNL1 levels in cell culture medium. Cells were stimulated by 10 µg/ml of poly(I:C). The concentration of supernatant cytokines was determined using multiplex MSD ELISA assay. Data are mean ±SD, n=3. *p<0.05, **p<0.005. **g.** The concentration of IFNL1 (left panel) and IFNL2 (right panel) in BeWo-Cas9 and -dA11 supernatants, measured by ELISA after exposure to 10 µg/ml of poly(I:C). Data are mean ±SD, n=6. ns = not significant, **p<0.01, ***p<0.001. **h**. Reporter gene activation by poly(I:C). Cells were transfected with luciferase reporters for IFNB, IFNL1, interferon-stimulated response element (ISRE) plus a Renilla-Luc control, then exposed to 10 μg/ml poly(I:C) overnight. Fold change was calculated *vs* unexposed BeWo-WT. Data are mean ±SD, *** p<0.001, n=3. **i**. A scatter plot depicting the correlation among poly(I:C)-induced ISG transcript in BeWo-A11 *vs* BeWo-WT (y-axis) and U2OS-WT *vs* U2OS-C19MC (x-axis). The graph shows mRNAs with ≥2-fold change and p<0.01 in both cell groups. The correlation coefficient (Pearson) is depicted in the graph.

To substantiate these findings, we used RT-qPCR to measure key ISGs at basal state and in response to poly(I:C) stimulation. As shown in Fig. 3d, we found a strong upregulation of selected ISGs in C19MC-deficient lines (dA11-21 and -23) in response to poly(I:C) when compared to BeWo-WT cells. As additional controls, we used BeWo cells that express Cas9, or two other BeWo-edited cells in which the deletion did not affect C19MC expression (clones BeWo-dA3/4, -21, and -41), thus ruling out the possibility that the elevated IFN response reflected lentivirus infection during genome editing. Using RT-qPCR, we also found increased levels of *IFNL*, the predominant IFN produced by trophoblast^22,33^, and to a lesser degree, *IFNB* RNAs (Fig. 3e). We confirmed our results by measuring the release of IFN proteins to cell culture media, using both Meso Scale Discovery (MSD) and ELISA assays (Fig. 3f-g). We also tested the effect of poly(I:C) on different IFN reporter systems, including ISRE-Luc, IFNB-Luc, and IFNL-Luc, using two separate BeWo-dA11 clones (BeWo-dA11-21 and -23). All three reporters exhibited activation upon poly(I:C) stimulation, with IFNL-Luc reporter consistently displaying the highest level of luciferase activity in BeWo-dA11 cells (Fig. 3h and Fig. S4a).

Using a proteomic analysis of BeWo cells exposed to poly(I:C), we validated the upregulation of ISGs among the top 100 proteins (Fig. S5). Lastly, we utilized our U2OS cell line, which we stably transfected with a BAC plasmid carrying the C19MC locus, enabling expression of the C19MC ncRNAs ^20^. Using RNA-seq to correlate expression changes between poly(I:C)- stimulated U2OS and BeWo cells that were engineered to express or to lack C19MC, we found a significant correlation between the two cell types, suggesting that C19MC expression in both systems led to the attenuated expression of innate immune genes (Fig 3i).

As PHT cells survive only several days in culture and thus cannot be *Cas9*-edited, we elected to recapitulate the effect of diminished C19MC expression on trophoblast stem cells (TS cells). We used H9-derived human TS cells that were differentiated into trophoblasts^34^, and edited the ATAC-11 element, using the same pair of single-guide RNAs (sgRNAs), as before. Our analysis confirmed the induction of ISGs in these TS cells (Fig 4a-c). Note that we used polyclonal TS cells and puromycin selection of the sgRNA-expressing construct and induction of expression using doxycycline. While these edited TS cells exhibited a very low proliferation rate^35^ that prevented us from establishing long-term selected clones, we were able to isolate several small TS cell clones and extracted sufficient RNA for RT-qPCR analysis of gene expression. These data recapitulated the phenotype of the batched TS cells (Fig. S6).

**Fig. 4:**
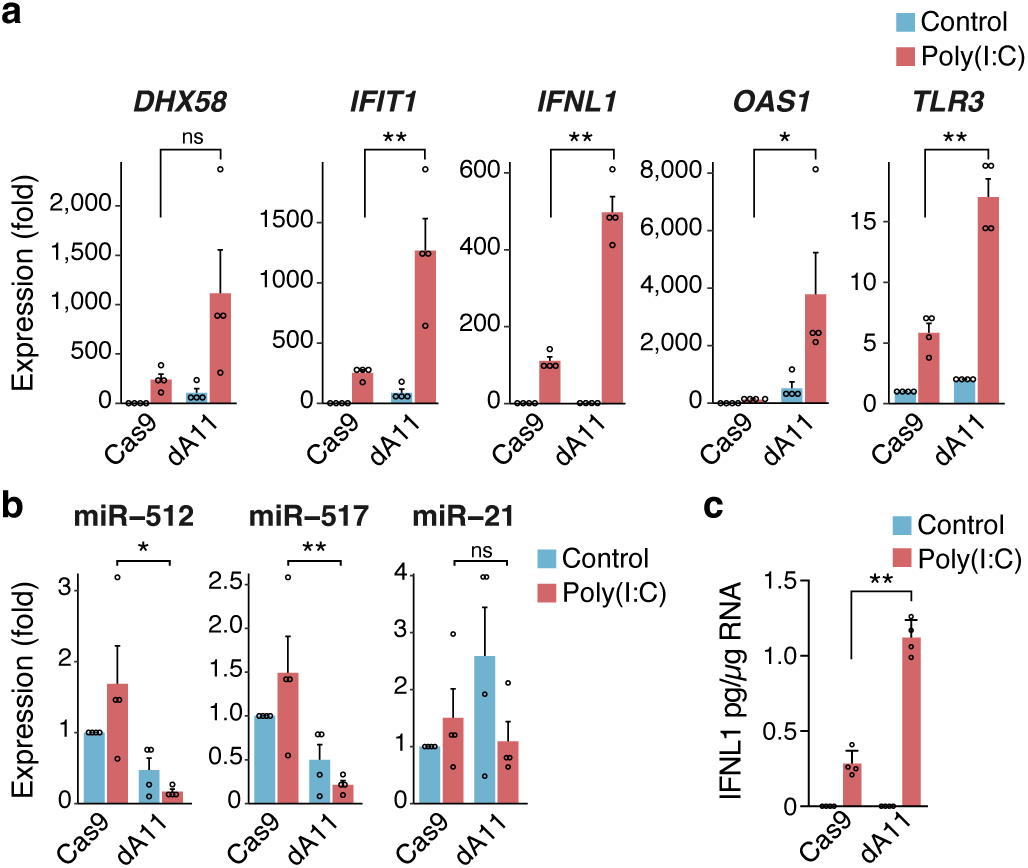
Interferon response C19MC-deleted human trophoblast stem cells (hTSCs). **a.** hTSCs were first transduced with a lentivector expressing Cas9 under the control of doxycycline, then with a lentivirus expressing two sgRNAs flanking the ATAC-11 element that was previously used to delete this element in BeWo cells. After selection with puromycin, polyclonal cells (dA11) were analyzed by RT-qPCR for selected ISGs and compared to control cells (Cas9). 10 µg/ml poly(I:C) was added overnight. Data are mean ±SD, n=3. ns = not significant, *p<0.05, **p<0.01. **b**. RT-qPCR analysis for selected miRNAs in control (Cas9) and edited polyclonal (dA11) hTSCs. 10 µg/ml poly(I:C) was added overnight. Data are mean ±SD, n=3. ns = not significant, *p<0.05, **p<0.01. **c**. IFNL1 levels in cell culture medium. Cells were unstimulated or stimulated with 10 µg/ml of poly(I:C) overnight. Supernatants were collected, and the concentration of IFNL1 was determined using ELISA. Data are mean ±SD, n=3. **p<0.01.

### Endosomal TLR3 mediates the augmented interferon response

As a key sensor for poly(I:C) is endosomal TLR3, which is expressed and functional in trophoblasts^36,37^, we compared the effect of poly(I:C) to that of agonists of other TLRs, including LPS (TLR4 agonist), imiquimod (TLR7/8 agonist), and ODN-2395 (TLR9 agonist). Using the ISG reporter assay, we found that only poly(I:C) elicited a strong activation of the reporters, supporting a role for TLR3 in the enhanced stimulation of ISGs in C19MC-mutant cells (Fig. 5a).

**Fig. 5:**
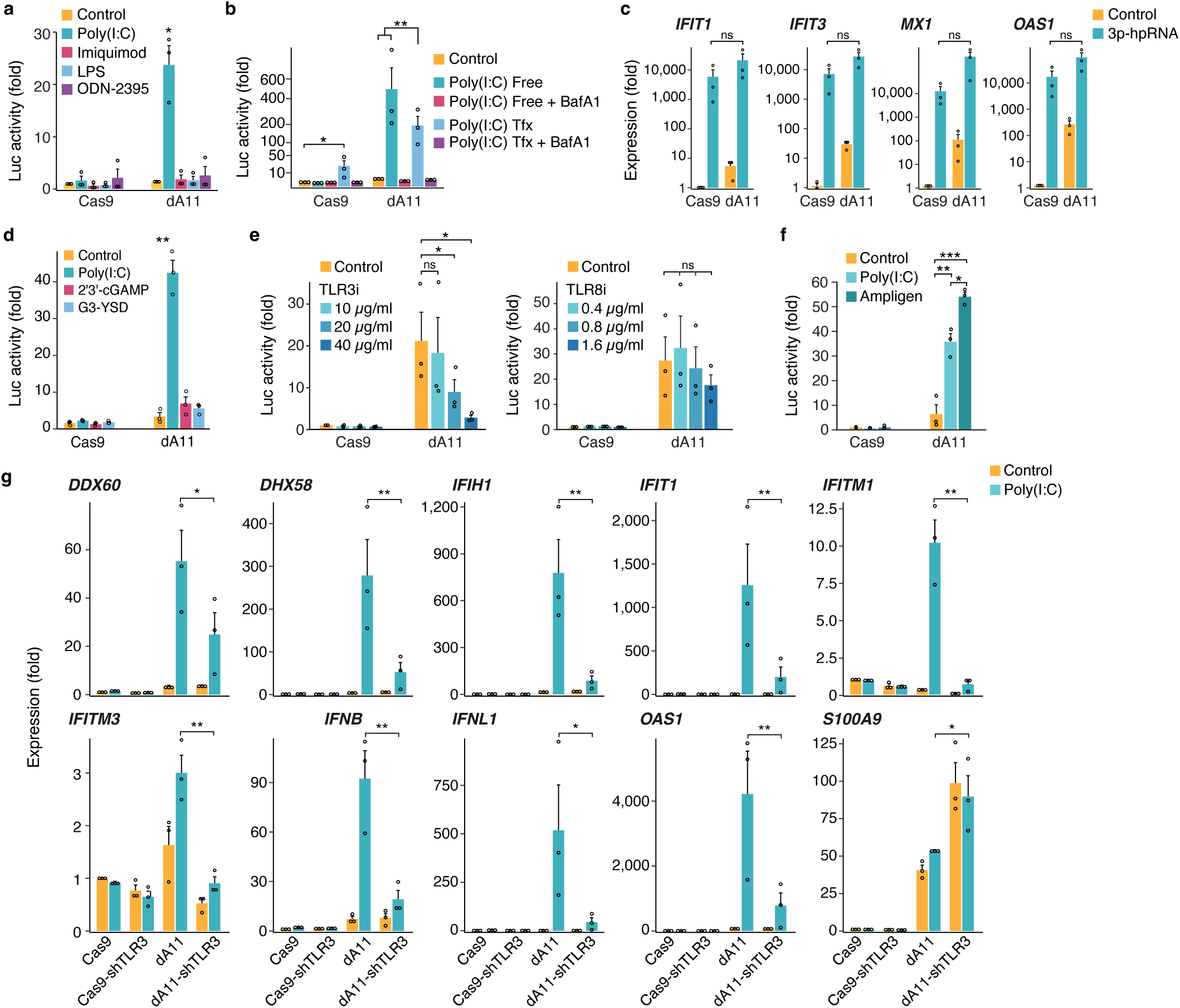
The influence of endosomal TLR3 on IFN response in C19MC-deleted cells. **a.** BeWo-Cas9 and -dA11 cells were transiently transfected with IFNB-luciferase (Luc) and a Renilla-Luc control (pRL-TK). The next day, the cells were exposed to various TLR ligands overnight. Fold change was calculated *vs* control BeWo-Cas9 cells. Bars are mean ±SD, n=3, each performed in duplicate. * p<0.001 compared to all other conditions. **b.** The role of endosomes. The cells were transiently as in a, and the next day, they were exposed to either free (floated) poly(I:C) or transfected with poly(I:C) using jetPRIME, in the presence or absence of bafilomycin A1. Bars are means ±SD, n=3, * p<0.01, ** p<0.001. **c.** The effect of 3p-hpRNA on ISG expression. Cells were transfected with 3p-hpRNA and incubated overnight. Total RNA was then isolated and analyzed by RT-qPCR. Bars are mean ±SD. n=3, each performed in duplicate. ns = nonsignificant. **d.** Activation of the cGAS-STING signaling in BeWo-Cas9 and - dA11 cells. Cells were transfected with IFNL1-luciferase reporter and exposed to poly(I:C), STING agonist (2’3’-cGAMP) and a cGAS agonist (G3-YSD) as described and analyzed in (a). **p<0.01. **e.** Inhibition of TLR3 or TLR8. BeWo-Cas9 and BeWo-dA11 cells expressing ISRE-Luc reporter were exposed to increasing concentration of a TLR3/dsRNA complex Inhibitor or a TLR8 inhibitor (CU-CPT9a) before being exposed to poly(I:C), 10 µg/ml, as described and analyzed in (a). * p<0.05. ns = nonsignificant. **f.** The effect of ampligen or poly(I:C), both at 10 µg/ml, on IFNB-Luc reporter activity, as described in (a). * p<0.05, ** p<0.005, *** p<0.0005. **g.** shRNA-mediated silencing of TLR3. Cells were transduced with a TLR3 shRNA lentivirus and selected for 10 days with puromycin. BeWo-Cas9 and -dA11 cells stably expressing shTLR3 were exposed to 10 µg/ml poly(I:C) before RT-qPCR. Data are mean ±SD. *p<0.05, ** p <0.01. n=3.

Poly(I:C) can also activate cytoplasmic sensors such as the helicases MDA5 and RIG-I^38,39^. To assess whether the activation of TLR3 in C19MC-deficient cells was mainly due to activation of endosomal TLR3, we compared the results of using (a) the addition of Poly(I:C) to cells known to activate mainly endosomal TLRs to (b) delivery of Poly(I:C) via transfection, known to also stimulate the cytoplasmic pathways^40,41^. When added directly to the cell cultures, we found that free poly(I:C) markedly stimulated the ISG reporter, but only in C19MC-depleted cells (Fig. 5b). In contrast, there was no difference between control and C19MC-deficient cells when poly(I:C) was delivered by transfection. Furthermore, the addition of bafilomycin A1, an inhibitor of endosomal acidification, prior to addition of free poly(I:C) completely blocked reporter activation, supporting a role for poly(I:C) in stimulating endosomal TLR3 (Fig. 5b and Fig. S4b). In contrast, 3p-hpRNA, which activates the RIG-I pathway, had a similar effect on control and dA11cells (Fig. 5c).

We also tested the effect of the DNA-sensing pathway. Using the cGAS agonist G3-YSD and the STING agonist 2’3’-cGAMP we detected an insignificant reporter activation by these agonists in BeWo-dA11 cells (Fig. 5d). Similarly, to rule out activation by double-stranded RNA (dsRNA)-sensing pathways, which might represent a response to immunogenic dsRNA from viruses, mitochondrial dsRNAs, or ALU elements (review in^42^), we stained the cells with an anti-dsRNA J2 antibody, and found a similar signal in control and dA11 mutant BeWo cells (Fig. S7). Similarly, inhibitors of the STING (H-151) and cGAS (G-140) pathways did not affect the reporter activity in the mutant cells (Fig. S8a). To further corroborate the role of TLR3 in our observed response to poly(I:C), we found that a specific inhibitor of TLR3 led to a concentration-dependent inhibition of the ISRE-luciferase reporter in BeWo-dA11 cells, while a TLR8 inhibitor exhibited a weak effect (Fig. 5e). Lastly, Ampligen, a specific TLR3 agonist used to treat several immune-mediated disorders ^43,44^, recapitulated the effect of poly(I:C) on C19MC-deficient BeWo cells, with no influence on control cells (Fig. 5f).

Our RNA-seq and RT-qPCR data indicated a higher expression of TLR3 in mutant cells even without an immune challenge, which we confirmed using western blot (Fig. S8B). To assess whether the enhanced TLR3 expression could account for the amplified innate immunity activation, we transfected BeWo-Cas9 cells with a plasmid expressing TLR3 (Fig. S8c) and tested for the effect of poly(I:C) stimulation on these cells. We found that overexpressing TLR3 led to a modest reporter induction in response to poly(I:C) stimulation (Fig. S8d), with no enhancement in the C19MC-depleted dA11 cells. Note that, in transfected BeWo-Cas9, the expression level of TLR3 was similar to that in the non-transfected BeWo-dA11 cells (Fig. S8c), suggesting that the enhanced IFN response in C19MC-depleted cells could not be attributed solely to elevated TLR3 expression. We further examined the role of TLR3 by knocking it down using a short hairpin RNA (shRNA) against TLR3. The downregulation of TLR3, which was confirmed by RT-qPCR (not shown) and western-blot (Fig. S8e), markedly attenuated the poly(I:C)-induced IFNL1-Luc reporter (Fig. S8f) and ISG expression in the BeWo-dA11 cells, with no effect in BeWo-Cas9 cells (Fig. 5g). Notably, as the TLR3 regulator S100A9^45^ is one of the top upregulated genes in our RNA-seq analysis of BeWo-sA11, we assessed whether the lower levels of S100A9 in BeWo-Cas9 cells limited their response to poly(I:C). We found that S100A9 overexpression did not alter the activation of the IFNL1-Luc reporter upon stimulation with poly(I:C) (Fig. S8g), suggesting that S100A9 expression does not play a major role in the heightened ISG response of BeWo-dA11 cells.

### Increased Poly(I:C) accumulation in the endosomal compartment of C19MC-depleted cells

Because small RNAs can compete with the uptake of poly(I:C) or its binding to TLR3^46,47^, we hypothesized that the high level of ncRNAs (C19MC miRNA or Alus) competes with the ability of poly(I:C) to bind endosomal TLR3. We first examined TLR3 localization in BeWo-Cas9 and - dA11 cells using three-dimensional confocal imaging. In BeWo-Cas9 cells, TLR3 was located in vesicles throughout the cytoplasm and frequently accumulated in the perinuclear area (Fig. S9a). Notably, in BeWo-dA11 cells, we observed a much stronger TLR3 immunofluorescence and even more pronounced concentration of TLR3-containing compartments in the perinuclear area when compared to that in BeWo-Cas9 (Fig. S9a). Interestingly, TLR3 was mainly located in LAMP1-positive vesicles (late endosomes and lysosomes) rather than in EEA1-positive early endosomes (for LAMP1, 20.13% in BeWo-Cas9 cells vs 35.84% of total cellular TLR3 in dA11 cells, and for EEA1, 4.96% in BeWo-Cas9 cells vs 11.78% of total cellular TLR3 in dA11 cells, p<0.001 for both comparisons, t-test), reflecting both the higher expression of TLR3 and a redistribution of its subcellular location. Under all conditions, the amount of TLR3 in early and late endosomes was significantly larger in BeWo-dA11 than BeWo-Cas9 cells (Fig S9b-c).

Next, using rhodamine-conjugated poly(I:C) (poly(I:C)-Rh), we detected significantly higher levels of poly(I:C) in TLR3-containing endosomes in BeWo-dA11 when compared to BeWo- Cas9 cells (Fig. 6a-b). To further support our findings, we tested the level of biotin-conjugated poly(I:C) in endosomes and found an increased retention of biotin-poly(I:C) in the endosomal fraction of BeWo-dA11 cells compared to BeWo-Cas9 cells, suggesting a more efficient binding of poly(I:C) to TLR3 in these cells (Fig. 6c).

**Fig. 6:**
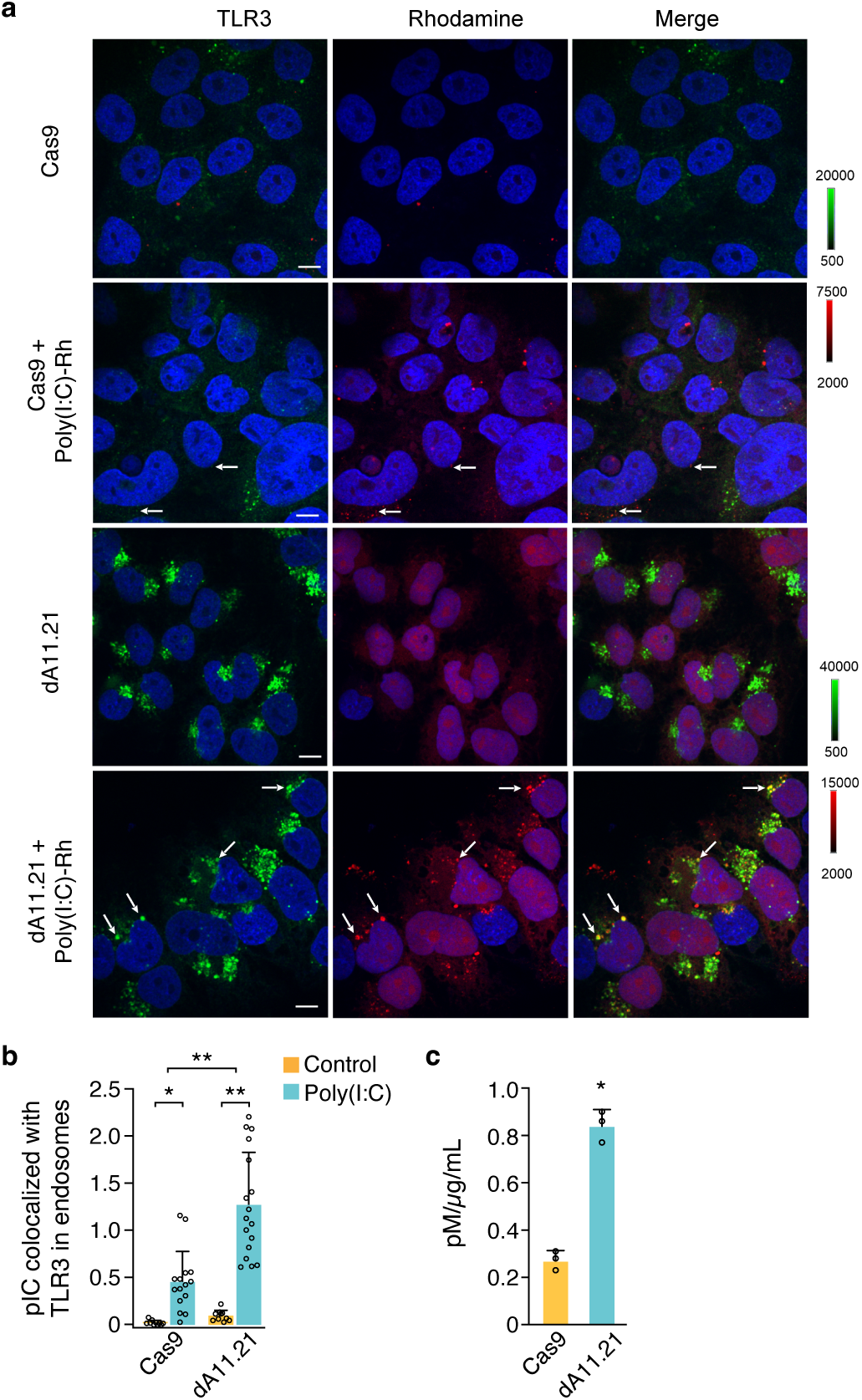
The colocalization poly(I:C) with TLR3. **a.** Single confocal sections of 3D images of BeWo-Cas9 or -dA11, exposed to poly(I:C)-Rh (10 µg/ml) overnight, then fixed and imaged for TLR3 immunofluorescence (488 nm, green), poly(I:C) (561 nm, red). DAPI staining was used for nuclei. The reddish color of the nuclei in BeWo-dA11 cells is due to the lentiviral vector used to deliver sgRNA, which also expresses mCherry. White arrows denote examples of colocalization of TLR3 and poly(I:C)-Rh. Note different fluorescence intensity scales due to higher levels of TLR3 and internalized poly(I:C)-Rh in BeWo-dA11 cells. Scale bars: 10 µm. **b**. Quantitative analysis of the amount of poly(I:C)-Rh colocalized with TLR3 in endosomes. Each bar represents ratio of poly(I:C)-Rh and TLR3 fluorescence in endosomes across multiple images exemplified in ***a***. Bars are mean ±SD. N=10-17 images, representing n=3. * p<0.05, **** p<0.0001. **c**. BeWo Cells were exposed to biotin-conjugated poly(I:C) (10 µg/ml) overnight, endosomes were isolated as described in Methods, and the biotin-poly(I:C) concentration was determined using a biotin quantification kit. Biotin concentration was normalized to the protein concentration in each sample. Data are mean ±SD. * p<0.005. n=3.

## Discussion

The balance between feto-placental and maternal immunity—preventing rejection of the semi-allogeneic fetus while responding to adverse microbes—remains a central question in the biology of human pregnancy and its influence on early life immune responses^48,49^. While interrogating the unique expression and function of C19MC RNAs, a primate- and placenta-specific, large miRNA cluster, we discovered that these small C19MC RNAs restrain the expression of key IFN-dependent components of trophoblastic innate immunity^22,33^. Unlike the immune stimulatory effect of C19MC small RNAs that we and others observed in *non-trophoblastic* cells ^20^ ^21^, here we observed a strong activation of IFN pathways in C19MC- depleted trophoblasts, which was accompanied by reduced viral replication in these cells.

It is unlikely that the robust ISG stimulation in C19MC-lacking trophoblasts is due to increased cellular dsRNA (and review in^42^) or, as recently reported, the effect of C19MC Alu short interspersed nuclear elements (SINEs) in *non-trophoblastic* HEK-293 cells. First, we markedly decreased, not increased, the total production of small ncRNAs. Second, we found no increase in total cellular RNA in the C19MC-deleted cells. Third, our results could not be explained by the lentiviral *cas9* transduction, as control cells were also transduced by several lentiviruses carrying single-guide RNAs targeting other regions of C19MC, but without impacting C19MC expression.

To abrogate the expression of C19MC RNAs, we first used ATAC-seq in primary trophoblasts and in trophoblast lines to delineate open chromatin sites in the C19MC 160 kb region, and validated the ATAC-seq-positive C19MC sites *in vivo* in transfected mouse trophectoderm.

Genomic ablation of the dd11 region, which is distinct from the known functional C19MC CpG island^25,35^, led to a marked (>1,000-fold) selective reduction in the expression of C19MC ncRNAs, with no effect on gene expression from other chromosome 19 regions, thus establishing that the C19MC A11 element is regulated in the human placenta in a cell-specific manner, likely by a trophoblast-specific transcriptional activator that is expressed from the first trimester, when C19MC is first detected in the plasma of pregnant women^15^.

Among pathogen-associated molecular patterns (PAMPs) that underlie virus interaction with host cell pattern recognition receptors (PRRs)^50^, we focused on TLR3, which responds to diverse RNA viruses through interaction of two TLR3 molecules with one dsRNA^51^ and translocates from its early endosomal location to late endosomes^45^. Activated TLR3 recruits TRIF to stimulate IRF3 and, subsequently, the production of IFNs and ISGs ^52,53^. Several lines of evidence implicate endosomal TLR3 in our observed enhancement of the trophoblastic IFN response: (a) the TLR3 agonist poly(I:C) markedly potentiated the observed innate immunity response, and this effect was recapitulated using several other TLR3 agonists and inhibited by a TLR3 dsRNA complex inhibitor or TLR3 knockdown; (b) the effect was strongest with direct application of poly(I:C) to cells, with a weaker effect when activating the cytoplasmic PRRs RIG-I and MDA5^54–56^;(c) bafilomycin, an inhibitor of endosomal acidification, abolished the ISG response; (d) G3-YSD, a GAS agonist, and 2’3’-cGAMP, a STING agonist, had virtually no effect in both WT and mutant cells. Together, our data position TLR3 as a key target for regulation by C19MC RNAs. Interestingly, the immune activating effect of Alu-SINEs in non-trophoblastic cells was TLR3 independent and included upregulation of IRF7 ^21^, further supporting a distinct effect of C19MC-depletion in trophoblasts.

How does the reduction in the level of C19MC ncRNAs potentiate ISG production? poly(I:C) is known to selectively activate TLR3 through entry via clathrin-mediated endocytosis, a process that is enhanced by CD14^57,58^. Small RNAs or double-stranded oligonucleotides can compete with either the uptake of poly(I:C) or its binding to TLR3 in endosomes^46,47,59^. Our results suggest that the abundance of C19MC RNAs might bind to TLR3 and block its activation by its endogenous or exogenous ligands. This is supported by (a) the difference between the amount of poly(I:C) with TLR3 in endosomes of WT and C19MC-mutant cells; (b) the difference in poly(I:C) levels in endosomes between the two cell types, and (c) [add competition experiment].

Across pregnancy, the immunological response to infections represents a balance of signals that originate from the maternal immune system and those from the fetal-placental unit. This balance reduces feto-placental infections while mitigating the risk of excessive immune activation, which has been implicated in a range of adverse neonatal outcomes^60,61^ (and reviewed in^62^). Our data suggest that C19MC RNAs dampen a potent immune response by placental trophoblasts, potentially instigated by dsRNAs that circulate in the maternal blood, and could affect key trophoblast functions and adversely impact pregnancy outcome (reviewed by^63^). Indeed, IFN-induced murine placental injury was found to be mitigated by IFNAR -/- knockout^64,65^. Further, discrete poly(I:C)-induced ISGs, such as IFITMs, have been implicated in decreased trophoblast fusion, placental abnormalities, and mouse resorption^66,67^.

Our results are limited by our inability to perform *Cas9*-deletion studies in PHT cells, as these senescent cells survive a very short time in culture and are not accessible to *Cas9* manipulations. Villous trophoblast cell lines that are known to express TLRs exhibited minimal ISG activation in response to their respective ligands. This includes TLR3 response to poly(I:C). In contrast, extravillous trophoblasts, which express very low levels of C19MC RNAs^27^, respond robustly to poly(I:C)^68^. We also do not know if there is active uptake and concentration of C19MC small RNAs in endosomal compartments, which may take place through breaches in the endosomal membrane lipid bilayer^69^ or by other unknown mechanisms.

In summary, innate immunity plays a critical role in feto-placental protection against microbial pathogens^70^. Yet, excessive innate or adaptive immune response might be detrimental for pregnancy health^71,72^. While small RNAs are known to regulate the immune system (reviewed in^73^), our data establishes an important, previously unrecognized homeostatic immune control mechanism in which C19MC-derived endogenous small ncRNAs guard human trophoblasts against overacting innate immunity.

## Methods

### Cell and culture

The Institutional Review Board at the University of Pittsburgh approved all placental procurement protocols that were used in these studies. PHT cells were isolated from healthy singleton term placentas under an exempt protocol approved by the University of Pittsburgh’s IRB (PRO08030033). Upon admission to the hospital, patients provided written informed consent for the use of de-identified and discarded tissues for research. PHT cells were dispersed from term placentas using a modification of previously published protocols^74,75^ and were maintained up to 72 h in Dulbecco Modified Eagle Medium (DMEM; Corning, On; New York, NY, USA) containing 10% bovine growth serum (BGS; HyClone, Logan, UT, USA) and antibiotics at 37°C in a 5% CO_2_ atmosphere. Human choriocarcinoma BeWo cells (ATCC, #CCL- 98, Manassas, VA, USA) were cultured in F-12K Kaighn’s modified medium (Corning), supplemented with GemCell SuperCalf serum (Gemini Bio-products, Sacramento, CA, USA) or Cosmic Calf serum (HyClone, Logan, UT, USA) and antibiotics. Immortalized human first trimester extravillous trophoblast cells (HTR-8/SVneo), provided by C.H. Graham, Kingston, Ontario, Canada^76^, were cultured in RPMI-1640 (CellGro, Manassas, VA, USA), supplemented with 5% GemCell SuperCalf serum (GeminiBio) and antibiotics. HeLa cells (ATCC, #CCL-2), U2OS (ATCC, #HTB-96), 293T (ATCC, #CRL-3216), HUVEC (ATCC, #PCS-100-010), were maintained in DMEM, supplemented with 10% GemCell SuperCalf serum (GeminiBio) and antibiotics. All cell lines were maintained in a humidified incubator at 37°C and 5% CO_2_. Human trophoblast stem cells (hTSCs, a gift of Thorold W. Theunissen, Washington University, St. Louis, MO, USA) were cultured as described^34^. Infection of BeWo cells with vesicular stomatitis virus (VSV, Indiana) was performed, using a multiplicity of infection (MOI) of 0.1 and 0.02 for 5.5 h, as previously described^20,77^.

### Assay for transposase-accessible chromatin using sequencing (ATAC-seq)

We performed ATAC-seq as described by Buenrostro *et al*.^26^, using PHT cells isolated from term placentas. HUVEC cells were used as reference because they are also primary cells but do not express C19MC miRNAs. Nuclei from 50,000 cells were isolated, and transposition was performed using transposase (Illumina, San Diego, CA, USA). Sequencing libraries were prepared using custom Nextera PCR primers (Illumina) as described by Buenrostro^26^. The ATAC-seq libraries for PHT and HUVEC cells were aligned to human reference genome assembly hg19 using Bowtie. To identify ATAC-seq peaks in our two libraries, we analyzed the data by ZINBA^78^, using a window size of 300 bp and an offset of 75 bp. Peaks were identified as those with a posterior probability > 0.99. Peak coordinates were converted to GRCh38 using liftOver (Bioconductor Package Maintainer (2024), *Changing genomic coordinate systems with rtracklayer::liftOver*. R package version 1.30).

### RNA sequencing (RNA-seq)

Total RNA samples were prepared from the cell lines, using TRI reagent (Molecular Research Center, Cincinnati, OH, USA) according to the manufacturer’s instructions, and purified using EconoSpin spin columns (Epoch Life Science, Missouri City, TX, USA) or PuroSpin NANO Silica Spin Columns (Luna Natotech, Markham, ON, Canada). An on-column DNase step, using RNase-free DNase (Qiagen, Valencia, CA, USA), was included. RNA libraries were prepared and sequenced at Novogene (Sacramento, CA, USA), using a NovaSeq PE150 platform. The RNA libraries were aligned to the human reference genome GRCh38, using the RNA-seq alignment tool STAR, and annotated with GENCODE (v. 27). The number of reads per mRNA/lncRNA was calculated for each RNA-seq library using STAR. For expression of Alu elements, we intersected the aligned bam files with annotation of Alu elements. An aligned read was considered as an Alu element if the length of the aligned read was at least 40 bp and at least 50% of its length overlapped with a known Alu element. Negative binomial test, as implemented in the R package DESeq2, was used to identify mRNA/lncRNAs, showing a significant change in expression between groups^79^. P-values were adjusted using Benjamini and Hochberg’s method to control the false discovery rate^80^.

### Transfection and reporter luciferase assay

The 19 open chromatin genomic regions identified by ATAC-seq were PCR-amplified from a BAC plasmid harboring the C19MC locus (RP11-1055O17)^24^ using primers containing restriction sites that were compatible with the pGL3p luciferase reporter (Promega, Madison, WI, USA). A list of the primers used for cloning is provided in Table S1. Constructs were verified by restriction and Sanger sequencing (Azenta Life Sciences, South Plainfield, NJ, USA). Using 12-well plates, each cell type was transfected with 1 µg of test sequences in pGL3p vector and 20 ng of Renilla luciferase control vector pGL4.74 (hRluc/TK, Promega), using polyethylenimine (PEI). PHT cells were transfected using lipofectamine 3000 (Thermo Fisher Scientific, Waltham, MA) according to the manufacturer’s instructions. Luciferase reporter plasmids for Renilla (Promega), p65-Luc, ISRE-Luc, and IFNB-Luc were previously described^20^. We also constructed an IFNL1-Luc in pGL3-basic reporter (Promega) by PCR amplification of a 2144-kb genomic DNA fragment upstream of the transcription start site (chr19:39,294,277-39,296,420, hg38), using primers IFNL1F and INFL1R (Table S1). BeWo cells, exposed to TLR agonists, were first transfected with individual luciferase reporters using PEI, and 24 h later, transfected with poly(I:C) (poly(I:C)-HMW, InvivoGen, San Diego, CA, USA) or 3p-hpRNA (InvivoGen), using jetPRIME (Polyplus, New York, NY, USA). Cells were incubated overnight, and luciferase activity was assayed using Dual-Luciferase Reporter Assay System (Promega). Firefly luciferase values were normalized to Renilla luciferase signals. Where indicated, cells were transfected with a construct expressing S100A9 (OriGene Technologies, Rockville, MD, USA).

### In vivo enhancer assay

To generate the lentiviral pLV-hsp68-eGFP, we obtained pLV-Enh eGFP Reporter-muKC2- IGR13 (Addgene, #59282, Watertown, MA, USA) and used XhoI and BamHI to replace the attB- enhancer with a double-strand oligo containing NheI, XbaI, and XhoI sites for cloning. The defined ATAC sequences were cloned in this vector upstream of the minimal mouse hsp68 promoter. The lentiviral vectors were transfected, using PEI, into 293T cells along with the envelope plasmid pMD2-G (Addgene, #12259) and the packaging plasmid psPAX2 (Addgene, #12260). The virus-containing culture media were harvested at 48 and 72 h, precleaned with a 500-g centrifugation and a 0.45-mm filtration (Corning, NY, USA), then overlaid on a 20% sucrose cushion in phosphate-buffered saline (PBS) and centrifuged at 11,000 rpm for 4 h. The viral titer was determined with a ZeptoMetrix p24 ELISA kit (ZeptoMetrix, Buffalo, NY, USA).

Trophoblast lineage-specific gene manipulations in mouse blastocysts were performed as described^81,82^. Briefly, blastocysts from superovulated females were harvested at E3.5 by flushing the uterine horns with FHM medium (MR-024-D, Millipore, Bellerica, MA, USA), and the zona pellucida was removed using acidic Tyrode’s solution (T1788, Sigma Aldrich, St. Louis, MO, USA). Zona-free blastocysts were incubated for 5 h with the lentivirus (∼1x10^7^ TU/mL) in KSOM medium, under mineral oil. Transduced blastocysts were transferred into E2.5 pseudo- pregnant mice. The feto-placental unit was harvested 14 days after embryo transfer (E17.5), and the placentas were imaged with a fluorescent dissecting microscope (Leica M80 equipped with a Lumencor Sola LED light engine (Leica Microsystems, Deerfield, IL, USA). All mouse experiments were approved by the Institutional Animal Care and Use Committee of the University of Pittsburgh (protocol 13092512) and conducted in accordance with United States Public Health Service Policy, as defined in the Guide for the Care and Use of Laboratory Animals prepared by the National Academy of Sciences.

### CRISPR/as9-based genomic editing

Paired single-guide RNA (sgRNA) sequences were designed by the CRISPETA tool (http://crispeta.crg.eu) and cloned into the pDECKO mCherry vector (Addgene, #78534)^83^.

BeWo cells stably expressing Cas9 under the control of doxycycline were produced by transduction with EdiT-R Inducible Lentiviral Cas9 (Horizon, Cambridge, UK). BeWo-Cas9 cells were transduced with pDECKO-sgRNAs, selected with 2 mg/ml puromycin 48 h after transduction, and grown in the presence of doxycycline (2 mg/ml) to induce the expression of Cas9. After 10 days, the cells were sorted to isolate single clones and PCR-screened using primers flanking the targeted region. The presence of the deletion in the edited region was confirmed by sequencing.

### Reverse transcription and quantitative PCR (RT-qPCR)

For analysis of cellular mRNA and miRNA, total RNA was extracted from mouse tissues and cell lines using TRI Reagent (Molecular Research Center), according to the manufacturer’s instructions, and purified using EconoSpin spin columns (Epoch) or PuroSpin NANO Silica Spin Columns (Luna Natotech, Markham, ON, Canada). An on-column DNase treatment of RNA samples using RNase-free DNase (Qiagen) was included in the protocol. RT-qPCR was performed in duplicate, using the ViiA 7 Sequence Detection System (Thermo Fisher) as previously described^14,29^ or QuantStudio 5 real-time PCR system (Thermo Fisher). For mRNA analysis, total RNA was reverse transcribed using the High-Capacity cDNA Reverse Transcription Kit (Thermo Fisher) according to the manufacturer’s protocol. Quantitative PCR was performed by means of SYBR Select (Thermo Fisher). For miRNA, cDNA synthesis and qPCR were performed with the miRScript PCR system or miRCURY LNA PCR system (both from Qiagen) according to the manufacturer’s protocols. PCR primers are shown in Table S1.

Dissociation curves were run on all reactions, and samples were normalized to the expression of YWHA or GAPDH for mRNAs and to RNU6B (Qiagen, cat #218300, GeneGlobe ID MS00033740), RNU5G (Qiagen, cat # 339306, GeneGlobe ID: YP00203909), or SNORD44 small nuclear RNAs ((Qiagen, cat # 339306, GeneGlobe ID: YP00203902) for miRNAs. The fold change relative to control samples was determined by the 2-ΔΔCt method^84^.

### RNA interference

A predesigned Mission shRNA-TLR3-GFP (clone TRCN0000056851, Sigma Aldrich) was used for TLR3 silencing. Lentiviral particles were generated by transfection of 293T cells with psPAX2 packaging (Addgene #12,260) and pMD2.G envelope plasmids (Addgene #12,259), using lipofectamine 3000 (Invitrogen/Thermo Fisher). BeWo-WT^Cas9^ and BeWo-dA11 cells were infected with virus supernatants in the presence of 8 mg/ml polybrene. GFP-positive cells expressing shTLR3 were then obtained by flow cytometric sorting.

### Western Immunoblotting

Cells were lysed on ice in RIPA buffer supplemented with Protease/Phosphatase Inhibitor Cocktail (Cell Signaling, Danvers, MA, USA) and sonicated three times, 7 sec each. Lysate protein concentrations were determined with Pierce BCA Protein Assay Kit (Thermo Fisher), using a VERSAmax microplate reader (Molecular Devices, San Jose, CA, USA). Protein samples (30 µg) were separated on SDS-PAGE and transferred onto a 0.2-µm polyvinylidene difluoride membrane (Bio-Rad, Hercules, CA, USA), using standard procedures. Membranes were immunoblotted with a mouse monoclonal TLR3 antibody (0.2 μg/ml, #AF1487, R&D Systems, Minneapolis, MN, USA). The blots were processed for chemiluminescence using the WesternBright Sirius kit (Advansta, San Jose, CA, USA).

### Immunofluorescence

BeWo cells were grown on glass coverslips in a 12-well plate and exposed to poly(I:C) (poly(I:C) HMW, InvivoGen) or poly(I:C)-Rh (InvivoGen) for the indicated time. The cells were fixed for 15 min with 4% formaldehyde in PBS and permeabilized for 15 min with 0.2% Triton X-100 in PBS. Coverslips were incubated for 1 h at room temperature with anti-TLR3 (4 μg/ml, mouse monoclonal, #40C1285.6 Novus Biologicals, Centennial, CO, USA), and in experiments with unlabeled Poly(I:C) (InvivoGen) with anti-EEA1 (1:50, rabbit monoclonal, Cell Signaling, Cat# 3288) or anti-LAMP1 (1:50, rabbit monoclonal, Cell Signaling, Cat#9091) in 2% bovine serum albumin (BSA) in PBS. The following secondary antibodies (4 μg/ml) were used: Goat anti-mouse IgG (H+L) Alexa fluor 488 (Thermo Fisher), Cy5 AffiniPure Donkey Anti-Rabbit IgG (H+L) (Jackson Laboratory, Bar Harbor, ME, USA). Coverslips were mounted with ProLong Gold Antifade Reagent with DAPI (Thermo Fisher).

Z-stacks of x-y confocal images were acquired using a Marianas spinning disk confocal imaging system based on Zeiss Axio Observer Z1 inverted fluorescence microscope system equipped with 63× Plan Apo PH NA1.4 objective and controlled by SlideBook software (Intelligent Imaging Innovation, Denver, CO, USA), as described^85^. All image acquisition settings were identical for all experimental variants in each experiment.

To quantitate the ratio of the amounts of poly(I:C)-Rh and TLR3 in endosomes, segment masks “TLR3” and “poly(I:C)” were generated from background-subtracted 3D images to select all voxels positive for TLR3 immunofluorescence (488 or 640 channels) and poly(I:C)-Rh fluorescence (561 nm channel). The ”Colocalization” mask, corresponding to voxels common in TLR3 and poly(I:C) masks, was generated. A “Nuclei” segment mask, corresponding to DAPI-positive voxels was also generated and subtracted from the Colocalization mask to eliminate voxels overlapping with the background 561-nm channel fluorescence of the nuclei to obtain the corrected Colocalization (“Colocalization-Corr”) mask. The sum of the fluorescence intensity of poly(I:C)-Rh, in arbitrary units, was divided by the sum of fluorescence intensity through the 488-nm channels (TLR3 immunofluorescence of the Colocalization-Corr mask to obtain an apparent ratio of polyIC-T poly(I:C)-Rh and TLR3 fluorescence in each field of view (FOV).

To quantitate the amounts of TLR3 in early and late endosomes in experiments with unlabeled poly(I:C), segment masks “TLR3” (488-nm channel) and “EEA1” or “LAMP1” (640-nm channel) were generated to select all voxels positive for TLR3 and EEA1 or LAMP1 immunofluorescence, respectively. A ”Colocalization” mask that corresponds to voxels common in TLR3 and EEA1 or LAMP1 masks, was generated. The sum of the fluorescence intensity of TLR3 was divided by the sum of fluorescence intensity of EEA1 or LAMP1 in the Colocalization mask to obtain an apparent ratio of TLR3 and EEA1 or LAMP1 immunofluorescence per field of view (FOV). To estimate the fraction of TLR3 in early and late endosomes of the total cellular TLR3, the sum of the fluorescence intensity of TLR3 in the Colocalization mask was divided by the sum of fluorescence intensity of the TLR3 mask per FOV.

DsRNA was detected using the J2 antibody (Anti-dsRNA Antibody, clone rJ2, Millipore-Sigma, Burlington, MA, USA) at 1/200 dilution in PBS with 0.5% BSA. The slides were incubated at 4°C overnight, washed and incubated with Alexa Fluor 488 goat anti-mouse at 4 mg/µl dilution. After mounting, the slides were viewed by confocal microscopy (Nikon A1R, Nikon Instruments, Melville, NY).

### Endosomal poly(I:C)-Biotin levels

Endosomes were isolated using the Minute Endosome Isolation and Cell Fractionation system (Invent Biotechnologies, # ED-028, Plymouth, MN, USA). Briefly, BeWo-Cas9 or -dA11 were exposed to poly(I:C)-Biotin (10 µg/ml) overnight. The cells were lysed as described for western immunoblotting and washed with cold PBS. Cell nuclei, membrane fragments, and larger organelles were sequentially removed, using the provided buffers and centrifugation as directed. Isolated endosomes were used for analysis of biotin levels, measured using the 0.1 ml QuantTag Biotin Quantification Kit (Vector Laboratories, Newark, CA, USA). Final biotin levels, normalized to endosomal total protein, were determined by spectrophotometry as described for for western immunoblotting.

### Cytogenetics analysis

These analyses were performed by the Cytogenetics and Cell Authentication Core at MD Anderson Cancer Center (Houston, TX). For chromosome instability, the cells were exposed to colcemid (0.04 µg/ml) overnight at 37°C and then to a hypotonic solution (0.075 M KCl) for 20 min at room temperature. The cells were fixed in methanol and acetic acid (3:1 by volume) for 15 min and washed three times in the fixative. The slides were air-dried and stained in 4% Giemsa stain. Thirty-five metaphases were analyzed from each cell line for chromosome aberrations, including breaks, fragments, fusions, ploidy, and chromosome number. Images were taken using 80i Nikon Microscope and Hiband Hyperspectral System Upgrade karyotyping workstations (Applied Spectral Imaging, Carlsbad, CA, USA). For FISH detection of the C19MC locus, a probe was derived from the BAC plasmid RP11-1055O17, detailed earlier, which was labeled with an aqua fluorophore. Two control probes, Spectrum Orange/19q13 and Spectrum Green 19p13 (Abbott, Abbott Park, IL, USA) were used for detecting nearby chromosome 19 regions.

### Gene ontology and pathway analysis

Gene ontology (GO) over-representation analysis on a set of genes, based on their GO annotations, was performed, using the clusterProfiler package implemented in R^86^. A dataset comprising the list of differentially expressed genes was analyzed using IPA software (Qiagen), performed with Core Analysis function in IPA, based on a fold change ≥ ±2, and a false discovery rate of p<0.05.

### Statistical analysis

All statistical analyses were performed with R software (R 4.3.2) or GraphPad Prism 10 software. Statistical significance for multiple comparison was calculated by one-way ANOVA, and Tukey post-hoc test or by two-tailed unpaired t-test, where appropriate. Values are presented as mean ±SD, derived from at least three independent experiments, as indicated in each figure legend. Significance was determined as p<0.05.

## Data availability

The RNAseq data were uploaded to Geo and assigned the accession number GSE293705.

## Acknowledgements

The project was supported by National Institutes of Health (NIH) grants R37HD086916 and R01HD103727, the Burkland Family Foundation, and the Twenty-Five Club of Magee-Womens Hospital (to Y.S.). We thank Lori Rideout for assistance in manuscript preparation and Bruce Campbell for editing.

## Author contributions

JFM and YS crafted the project, designed the experiments and wrote the manuscript. JFM, YO, ES, VKK, HLS, LJB, SNS, and YS carried out the experiments and analyzed the data. AS designed and performed confocal imaging experiments, analyzed the images and related data. TC conducted the analysis of high-throughput data and statistical analysis. YS acquired funding and supervised the project.

## Competing interests

Y.S. is a consultant to Bio-Rad Laboratories, Inc. The authors certify that there are no financial interests that influenced the results and findings obtained through this study.

## Notes

### Summary of Updates

Corrections were made in the Methods (page 16, lines 5-9 and page 19, lines 20-24) to confirm details of the institutional approval for human subjects and mouse research, and reference 2 (page 26), which was incomplete, has been corrected.

## References

1. Turco, M. Y. & Moffett, A. Development of the human placenta. Development 146, (2019).

2. Sadovsky, Y. & Jansson, T. in Knobil and Neill’s Physiology of Reproduction (4th Edition) (eds T.M. Plant & A.J. Zeleznik) 1741–1782 (Academic Press, 2015).

3. Morelli, A. E. & Sadovsky, Y. Extracellular vesicles and immune response during pregnancy: A balancing act. Immunological reviews 308, 105–122, (2022).

4. Brosens, I., Pijnenborg, R., Vercruysse, L. & Romero, R. The "Great Obstetrical Syndromes" are associated with disorders of deep placentation. Am J Obstet Gynecol 204, 193–201, (2011).

5. Hemberger, M., Hanna, C. W. & Dean, W. Mechanisms of early placental development in mouse and humans. Nat Rev Genet 21, 27–43, (2020).

6. Lee, C. Q. et al. What Is Trophoblast? A Combination of Criteria Define Human First-Trimester Trophoblast. Stem Cell Reports 6, 257–272, (2016).

7. Muhlhauser, J., Crescimanno, C., Kasper, M., Zaccheo, D. & Castellucci, M. Differentiation of human trophoblast populations involves alterations in cytokeratin patterns. J Histochem Cytochem 43, 579–589, (1995).

8. Hemberger, M., Udayashankar, R., Tesar, P., Moore, H. & Burton, G. J. ELF5-enforced transcriptional networks define an epigenetically regulated trophoblast stem cell compartment in the human placenta. Hum Mol Genet 19, 2456–2467, (2010).

9. Apps, R. et al. Human leucocyte antigen (HLA) expression of primary trophoblast cells and placental cell lines, determined using single antigen beads to characterize allotype specificities of anti-HLA antibodies. Immunology 127, 26–39, (2009).

10. Bortolin-Cavaille, M. L., Dance, M., Weber, M. & Cavaille, J. C19MC microRNAs are processed from introns of large Pol-II, non-protein-coding transcripts. Nucleic Acids Res 37, 3464–3473, (2009).

11. Bentwich, I. A postulated role for microRNA in cellular differentiation. FASEB J 19, 875–879, (2005).

12. Chen, L. L. & Yang, L. ALUternative Regulation for Gene Expression. Trends Cell Biol 27, 480–490, (2017).

13. Donker, R. B. et al. The expression profile of C19MC microRNAs in primary human trophoblast cells and exosomes. Mol Hum Reprod 18, 417–424, (2012).

14. Ouyang, Y., et al. Term Human Placental Trophoblasts Express SARS-CoV-2 Entry Factors ACE2, TMPRSS2, and Furin. mSphere 6, (2021).

15. Dumont, T. M. F. et al. The expression level of C19MC miRNAs in early pregnancy and in response to viral infection. Placenta 53, 23–29, (2017).

16. Li, M. et al. Frequent amplification of a chr19q13.41 microRNA polycistron in aggressive primitive neuroectodermal brain tumors. Cancer Cell 16, 533–546, (2009).

17. Raghuram, N., Khan, S., Mumal, I., Bouffet, E. & Huang, A. Embryonal tumors with multi-layered rosettes: a disease of dysregulated miRNAs. J Neurooncol 150, 63–73, (2020).

18. Fonseca, A. Y. G., Santos, J. G. & Aristizabal-Pachon, A. F. An Overview of C19MC Cluster Subgroup 3 and Cancer. Microrna 10, 154–163, (2021).

19. Flor, I. & Bullerdiek, J. The dark side of a success story: microRNAs of the C19MC cluster in human tumours. J Pathol 227, 270–274, (2012).

20. Delorme-Axford, E. et al. Human placental trophoblasts confer viral resistance to recipient cells. Proc Natl Acad Sci U S A 110, 12048–12053, (2013).

21. Wickramage, I. et al. SINE RNA of the imprinted miRNA clusters mediates constitutive type III interferon expression and antiviral protection in hemochorial placentas. Cell host & microbe 31, 1185–1199 e1110, (2023).

22. Bayer, A. et al. Type III Interferons Produced by Human Placental Trophoblasts Confer Protection against Zika Virus Infection. Cell host & microbe 19, 705–712, (2016).

23. Chang, G. et al. Expression and trafficking of placental microRNAs at the feto-maternal interface. FASEB J, (2017).

24. Mouillet, J. F. et al. Transgenic expression of human C19MC miRNAs impacts placental morphogenesis. Placenta 101, 208–214, (2020).

25. Noguer-Dance, M. et al. The primate-specific microRNA gene cluster (C19MC) is imprinted in the placenta. Hum Mol Genet 19, 3566–3582, (2010).

26. Buenrostro, J. D., Giresi, P. G., Zaba, L. C., Chang, H. Y. & Greenleaf, W. J. Transposition of native chromatin for fast and sensitive epigenomic profiling of open chromatin, DNA-binding proteins and nucleosome position. Nat Methods 10, 1213–1218, (2013).

27. Xie, L. et al. C19MC microRNAs regulate the migration of human trophoblasts. Endocrinology 155, 4975–4985, (2014).

28. Nguyen, T. A. et al. High-throughput functional comparison of promoter and enhancer activities. Genome Res 26, 1023–1033, (2016).

29. Mishima, T., Miner, J. H., Morizane, M., Stahl, A. & Sadovsky, Y. The expression and function of fatty acid transport protein-2 and -4 in the murine placenta. PLoS One 6, e25865, (2011).

30. Pattillo, R. A. & Gey, G. O. The establishment of a cell line of human hormone-synthesizing trophoblastic cells in vitro. Cancer Res 28, 1231–1236, (1968).

31. Xie, K., Minkenberg, B. & Yang, Y. Boosting CRISPR/Cas9 multiplex editing capability with the endogenous tRNA-processing system. Proc Natl Acad Sci U S A 112, 3570–3575, (2015).

32. Matsuda, T. & Cepko, C. L. Electroporation and RNA interference in the rodent retina in vivo and in vitro. Proc Natl Acad Sci U S A 101, 16–22, (2004).

33. Wells, A. I. & Coyne, C. B. Type III Interferons in Antiviral Defenses at Barrier Surfaces. Trends Immunol 39, 848–858, (2018).

34. Dong, C. et al. Derivation of trophoblast stem cells from naive human pluripotent stem cells. Elife 9, (2020).

35. Kobayashi, N. et al. The microRNA cluster C19MC confers differentiation potential into trophoblast lineages upon human pluripotent stem cells. Nature communications 13, 3071, (2022).

36. Pineda, A., Verdin-Teran, S. L., Camacho, A. & Moreno-Fierros, L. Expression of toll-like receptor TLR-2, TLR-3, TLR-4 and TLR-9 is increased in placentas from patients with preeclampsia. Arch Med Res 42, 382–391, (2011).

37. Motomura, K. et al. Comprehensive Analysis of the Expression and Functions of Pattern Recognition Receptors in Differentiated Cytotrophoblasts Derived from Term Human Placentas. J Immunol 210, 1552–1563, (2023).

38. Kato, H. et al. Cell type-specific involvement of RIG-I in antiviral response. Immunity 23, 19–28, (2005).

39. Kato, H. et al. Differential roles of MDA5 and RIG-I helicases in the recognition of RNA viruses. Nature 441, 101–105, (2006).

40. Dauletbaev, N., Cammisano, M., Herscovitch, K. & Lands, L. C. Stimulation of the RIG- I/MAVS Pathway by Polyinosinic:Polycytidylic Acid Upregulates IFN-beta in Airway Epithelial Cells with Minimal Costimulation of IL-8. J Immunol 195, 2829–2841, (2015).

41. Palchetti, S. et al. Transfected poly(I:C) activates different dsRNA receptors, leading to apoptosis or immunoadjuvant response in androgen-independent prostate cancer cells. J Biol Chem 290, 5470–5483, (2015).

42. Hur, S. Double-Stranded RNA Sensors and Modulators in Innate Immunity. Annu Rev Immunol 37, 349–375, (2019).

43. Jasani, B., Navabi, H. & Adams, M. Ampligen: a potential toll-like 3 receptor adjuvant for immunotherapy of cancer. Vaccine 27, 3401–3404, (2009).

44. Navabi, H. et al. A clinical grade poly I:C-analogue (Ampligen) promotes optimal DC maturation and Th1-type T cell responses of healthy donors and cancer patients in vitro. Vaccine 27, 107–115, (2009).

45. Tsai, S. Y. et al. Regulation of TLR3 Activation by S100A9. J Immunol 195, 4426–4437, (2015).

46. Leonard, J. N. et al. The TLR3 signaling complex forms by cooperative receptor dimerization. Proc Natl Acad Sci U S A 105, 258–263, (2008).

47. Skold, A. E. et al. Single-stranded DNA oligonucleotides inhibit TLR3-mediated responses in human monocyte-derived dendritic cells and in vivo in cynomolgus macaques. Blood 120, 768–777, (2012).

48. Mor, G., Aldo, P. & Alvero, A. B. The unique immunological and microbial aspects of pregnancy. Nat Rev Immunol 17, 469–482, (2017).

49. Lim, A. I. et al. Prenatal maternal infection promotes tissue-specific immunity and inflammation in offspring. Science 373, (2021).

50. Kawai, T. & Akira, S. The role of pattern-recognition receptors in innate immunity: update on Toll-like receptors. Nat Immunol 11, 373–384, (2010).

51. Luo, J. et al. Lateral clustering of TLR3:dsRNA signaling units revealed by TLR3ecd:3Fabs quaternary structure. J Mol Biol 421, 112–124, (2012).

52. Barton, G. M. & Kagan, J. C. A cell biological view of Toll-like receptor function: regulation through compartmentalization. Nat Rev Immunol 9, 535–542, (2009).

53. Lee, B. L. & Barton, G. M. Trafficking of endosomal Toll-like receptors. Trends Cell Biol 24, 360–369, (2014).

54. Chow, K. T., Gale, M., Jr. & Loo, Y. M. RIG-I and Other RNA Sensors in Antiviral Immunity. Annu Rev Immunol 36, 667–694, (2018).

55. Vazquez, C. & Horner, S. M. MAVS Coordination of Antiviral Innate Immunity. J Virol 89, 6974–6977, (2015).

56. Akira, S., Uematsu, S. & Takeuchi, O. Pathogen recognition and innate immunity. Cell 124, 783–801, (2006).

57. Itoh, K., Watanabe, A., Funami, K., Seya, T. & Matsumoto, M. The clathrin-mediated endocytic pathway participates in dsRNA-induced IFN-beta production. J Immunol 181, 5522–5529, (2008).

58. Lee, H. K., Dunzendorfer, S., Soldau, K. & Tobias, P. S. Double-stranded RNA-mediated TLR3 activation is enhanced by CD14. Immunity 24, 153–163, (2006).

59. Liu, L. et al. Structural basis of toll-like receptor 3 signaling with double-stranded RNA. Science 320, 379–381, (2008).

60. Meuwissen, M. E. et al. Human USP18 deficiency underlies type 1 interferonopathy leading to severe pseudo-TORCH syndrome. J Exp Med 213, 1163–1174, (2016).

61. Cappelletti, M., et al. Type I interferons regulate susceptibility to inflammation-induced preterm birth. JCI Insight 2, e91288, (2017).

62. Ding, J. et al. Mechanisms of immune regulation by the placenta: Role of type I interferon and interferon-stimulated genes signaling during pregnancy. Immunological reviews 308, 9–24, (2022).

63. Yockey, L. J. & Iwasaki, A. Interferons and Proinflammatory Cytokines in Pregnancy and Fetal Development. Immunity 49, 397–412, (2018).

64. Miner, J. J. et al. Zika Virus Infection during Pregnancy in Mice Causes Placental Damage and Fetal Demise. Cell 165, 1081–1091, (2016).

65. Yockey, L. J., et al. Type I interferons instigate fetal demise after Zika virus infection. Sci Immunol 3, (2018).

66. Buchrieser, J. et al. IFITM proteins inhibit placental syncytiotrophoblast formation and promote fetal demise. Science 365, 176–180, (2019).

67. Degrelle, S. A., et al. IFITM1 inhibits trophoblast invasion and is induced in placentas associated with IFN-mediated pregnancy diseases. iScience 26, 107147, (2023).

68. Gierman, L. M. et al. Toll-like receptor profiling of seven trophoblast cell lines warrants caution for translation to primary trophoblasts. Placenta 36, 1246–1253, (2015).

69. Dowdy, S. F. Endosomal escape of RNA therapeutics: How do we solve this rate-limiting problem? RNA 29, 396–401, (2023).

70. Ander, S. E., Diamond, M. S. & Coyne, C. B. Immune responses at the maternal-fetal interface. Sci Immunol 4, (2019).

71. Chavan, A. R., Griffith, O. W. & Wagner, G. P. The inflammation paradox in the evolution of mammalian pregnancy: turning a foe into a friend. Curr Opin Genet Dev 47, 24–32, (2017).

72. Moffett, A. & Loke, C. Immunology of placentation in eutherian mammals. Nat Rev Immunol 6, 584–594, (2006).

73. Mehta, A. & Baltimore, D. MicroRNAs as regulatory elements in immune system logic. Nat Rev Immunol 16, 279–294, (2016).

74. Kliman, H. J., Nestler, J. E., Sermasi, E., Sanger, J. M. & Strauss, J. F., 3rd. Purification, characterization, and in vitro differentiation of cytotrophoblasts from human term placentae. Endocrinology 118, 1567–1582, (1986).

75. Nelson, D. M., Johnson, R. D., Smith, S. D., Anteby, E. Y. & Sadovsky, Y. Hypoxia limits differentiation and up-regulates expression and activity of prostaglandin H synthase 2 in cultured trophoblast from term human placenta. Am J Obstet Gynecol 180, 896–902, (1999).

76. Graham, C. H. et al. Establishment and characterization of first trimester human trophoblast cells with extended lifespan. Exp Cell Res 206, 204–211, (1993).

77. Bayer, A. et al. Chromosome 19 microRNAs exert antiviral activity independent from type III interferon signaling. Placenta 61, 33–38, (2018).

78. Rashid, N. U., Giresi, P. G., Ibrahim, J. G., Sun, W. & Lieb, J. D. ZINBA integrates local covariates with DNA-seq data to identify broad and narrow regions of enrichment, even within amplified genomic regions. Genome Biol 12, R67, (2011).

79. Love, M. I., Huber, W. & Anders, S. Moderated estimation of fold change and dispersion for RNA-seq data with DESeq2. Genome Biol 15, 550, (2014).

80. Benjamini, Y. Controlling the False Discovery Rate: A Practical and Powerful Approach to Multiple Testing. Journal of the Royal Statistical Society. Series B (Methodological) 57, 289–300, (1995).

81. Okada, Y. et al. Complementation of placental defects and embryonic lethality by trophoblast-specific lentiviral gene transfer. Nat Biotechnol 25, 233–237, (2007).

82. Mishima, T., Sadovsky, E., Gegick, M. E. & Sadovsky, Y. Determinants of effective lentivirus-driven microRNA expression in vivo. Scientific reports 6, 33345, (2016).

83. Aparicio-Prat, E. et al. DECKO: Single-oligo, dual-CRISPR deletion of genomic elements including long non-coding RNAs. BMC Genomics 16, 846, (2015).

84. Livak, K. J. & Schmittgen, T. D. Analysis of relative gene expression data using real-time quantitative PCR and the 2(-Delta Delta C(T)) Method. Methods 25, 402–408, (2001).

85. Sorkina, T. et al. Monoamine transporter ubiquitination and inward-open conformation synergistically maximize transporter endocytosis. Sci Adv 10, eadq9793, (2024).

86. Yu, G., Wang, L. G., Han, Y. & He, Q. Y. clusterProfiler: an R package for comparing biological themes among gene clusters. OMICS 16, 284–287, (2012).

