## Supplemental Figures 1-9 and Supplemental Table 1 for "The Chromosome 19 miRNA Cluster Guards Trophoblasts Against Overacting Innate Immunity"

#### Supplementary Figures

**Fig S1. Analysis of the C19MC enhancer elements**. **a**. Luciferase (Luc) activity of "short" ATAC elements versus "long" element in BeWo cells (Fig. 1). The short elements corresponded to sequences identified by ATAC-seq, ranging from 133 bp (ATAC-12) to 751 bp (ATAC-11). The long elements included these short sequences, with additional flanking regions, extending the total length to approximately 2 kb. These sequences were cloned in the pGL3-reporter plasmid and transiently transfected, along with a *Renilla* Luc plasmid, for normalization. Data are mean ±SD (n=5). ns, nonsignificant. **b**. Sequences corresponding to the four functional ATAC peaks were cloned in the pGL3-promoter reporter plasmid in sense and antisense orientation and transiently transfected as in (a). Luc activity for each ATAC region was compared to the empty pGL3-promoter plasmid. Data are mean ±SD. ns: nonsignificant. n=3. **c**. C19MC RNA output in the various ATAC-element deleted cells. RT-qPCR analysis of miR-517 expression in BeWo cells across different clones for each ATAC-3/4, ATAC-12, and ATAC-17 edited elements. Multiple independent clones for each deletion were tested for miR-517a expression and compared to wild-type (WT) cells and an ATAC-11 edited clone. Each graph represents a single experiment.

**Fig S2. FISH validation of chromosome 19 copy number. a**. FISH analysis of mitotic chromosomal spreads in BeWo-Cas9 and BeWo-dA11 cells, using a C19MC probe (aqua blue)

and two control probes for c19q13 and c19p13 (green). Both cell lines exhibit triploidy for the C19MC locus. The lower insets represent an enlargement of the square box in the larger pictures. **b**. Chromosome counts from cytogenetic analysis performed on 20 chromosome spread samples in BeWo-Cas9 and BeWo-dA11 cells. **c**. The results of chromosome instability analysis conducted on 35 metaphase chromosome spreads for each cell line.

**Fig S3: C19MC-altered signaling pathways. a:** Enrichment analysis, illustrating GO biological processes that are associated with abrogated C19MC expression in trophoblasts. The figure shows the top eight significantly regulated GO biological processes. **b**. A graphical summary of IPA core analysis of differentially expressed genes in BeWo-Cas9 *vs* -dA11 cells.

**Fig S4. An additional clone of C19MC-silenced cells (BeWo-dA11.23) and innate immune response.** The responses recapitulate the findings using the BeWo-dA11 line. **a.** Activation of various Luciferase (Luc) reporter genes by poly(I:C). The cells were transfected with a Luc reporter driven by the interferon-β promoter (IFNB), interferon-λ1 promoter (IFNL1), interferon-stimulated response element (ISRE), along with a Renilla-Luc internal control. The next day, cells were exposed to poly(I:C) (10 μg/ml) overnight before the Luc assay. The fold change was calculated relative to the mean of BeWo-WT cells without poly(I:C). Data are mean ±SD, n=3. ** p<0.001. **b**. RT-qPCR analysis of interferon-λ1 (*IFNL1*), interferon-λ2 (*IFNL2*), interferon-β (*IFNB*) and *OAS1* in BeWo-dA11 clone 23. 10 μg/ml poly(I:C) was added overnight. Data are mean ±SD, n=3. ** p<0.001.

**Fig S5: A proteomics analysis, focusing on expressed genes identified by transcriptomics.** The graph displays the 100 most upregulated or downregulated transcripts in BeWo-dA11 cells vs BeWo-Cas9 cells, matched with the 100 most upregulated or downregulated proteins. Genes present in both analyses are red-shaded, while those found only by transcriptomics are grey-shaded.

**Fig. S6: An innate immune response in the cloned lines of C19MC-dA11-deleted hTSCs**. Each of the three panels represents the data from an independently edited clone that survived long enough in culture to yield sufficient RNA for RT-PCR analysis of a small set of genes. For each clone, the main panel shows INFL1 or OAS1 expression in edited dA11 *vs* Cas9 control cells, after poly(I:C) stimulation. The inset panels show miR-517 expression in control vs dA11-edited cells exposed to 10 µg/ml poly(I:C) before RT-qPCR.

**Fig S7: Immunofluorescence analysis of BeWo cells for the presence of dsRNA**. Detection of dsRNA in BeWo cells. BeWo-Ctr and BeWo-dA11 were stained with the anti-dsRNA antibody (J2, green signal) and DAPI (blue). Cells were then observed under a laser confocal microscope. Scale bar =50 µm. The images are representative of at least 3 independent experiments.

**Fig. S8**: **TLR3-mediated innate immune response in C19MC-deficient BeWo cells. a.** Inhibitors of the cGAS-STING pathway do not alter the interferon response in BeWo cells. Cells were transfected with the IFNL1- luciferase (Luc) reporter and exposed to H-151 (10 µM) or G-140 (10 µM) for 2 h. Poly(I:C) 10 µg/ml was added overnight. Data are mean ±SD, n=3. ns = not significant. **b**. Western blot analysis of TLR3 in BeWo-Cas9 and -dA11 cells. The cells were stimulated with poly(I:C) (10 µg/ml) for 6 h and analyzed by western blot, as described in Methods, using an anti-TLR3 antibody. Actin was used as a loading control. **c**. Overexpression of TLR3. BeWo-Cas9 and -dA11 cells were transiently transfected with a TLR3 expression plasmid for 48 h, then analyzed by western blot, using a TLR3 antibody. Actin was used as a loading control. **d**. BeWo-Cas9 and -dA11 cells were transfected with IFNL1-Luc reporter and a plasmid expressing TLR3, as described in Methods. The next day, cells were incubated with 10 μg/ml poly(I:C). Data are mean ±SD, **p<0.005. n=3. **e**. shRNA-mediated silencing of TLR3. Cells were transduced with a TLR3 shRNA lentivirus and selected for 10 days with puromycin. Western blot analysis of TLR3 protein from cell lysates of BeWo-Cas9 and BeWo-dA11. Tubulin was used as a loading control. **f**. BeWo-Cas9 and -dA11 cells expressing shTLR3 were transiently transfected with IFNL1-Luc reporter. The next day, cells were incubated with 10 μg/ml poly(I:C) for 6 h. Data are mean ±SD, **p<0.001. n=5. **g**. BeWo-Cas9 and -dA11 cells were transiently transfected with IFNL1-Luc reporter and a construct expressing S100A9, as described in Methods. The next day, cells were incubated with 10 μg/ml poly(I:C). Data are mean ±SD, ns, not significant. n=3. The inset RT-qPCR analysis (a single experiment) confirms S100A9 overexpression in cells transfected with the S100A9 construct or a control plasmid (pBluescript). The graph represents data from a single RT-qPCR experiment.

**Fig. S9. Localization of TLR3 in early and late endosomes. a.** Single confocal sections of 3D images of BeWo-Cas9 and BeWo-dA11 exposed to poly(I:C) (10 μg/ml) for 2 h or 16 h, then fixed, immunolabeled and imaged for TLR3 (488 nm, green), EEA1 (640 nm, red) or LAMP1 (640 nm, red). DAPI staining (blue) was used for nuclei. Fluorescent intensity scales are identical in all images. Scale bars: 10 μm. **b and c**. Quantitative analysis of the amount of poly(I:C)-Rh colocalized with TLR3 in endosomes. Each bar represents ratio of the TLR3 fluorescence to EEA1 (early endosomes) or LAMP1 fluorescence (late endosomes and lysosomes) across multiple 3D images exemplified in ***a***. Bars are mean ±SD. Images represent n=5. * p<0.0005 for the left panel, and ** p<0.0001 for the right panel.

### Supplementary Figure 1

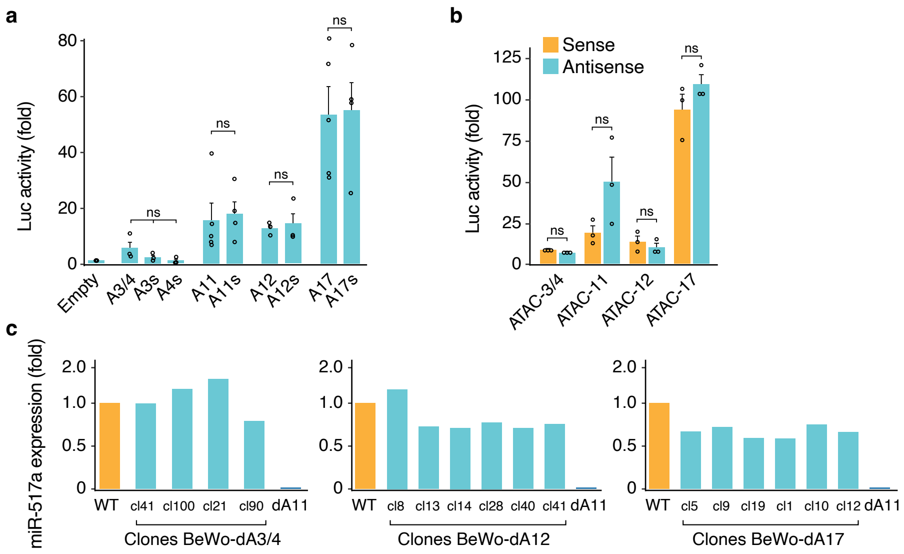

### Supplementary Figure 2

**a**

Cas9 dA11

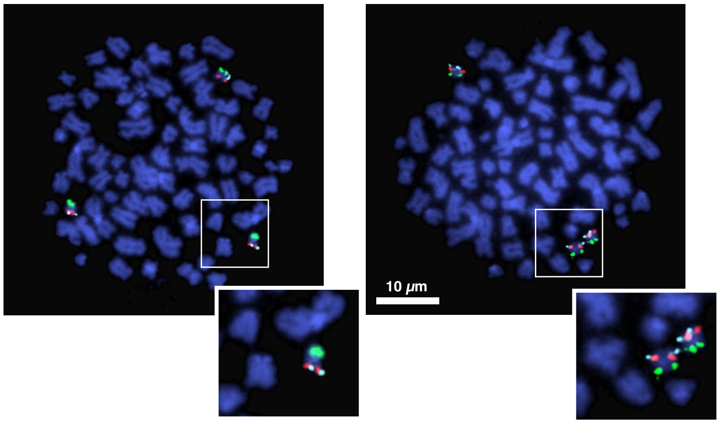


**b**

**Suppl. Table 1.** Chromosome counts.

| | BeWo Cas9 | BeWo dA11 |
|---|---|---|
| 1 | 77 | 71 |
| 2 | 75 | 75 |
| 3 | 64 | 78 |
| 4 | 68 | 73 |
| 5 | 75 | 71 |
| 6 | 74 | 72 |
| 7 | 77 | 71 |
| 8 | 78 | 63 |
| 9 | 77 | 141 |
| 10 | 77 | 67 |
| 11 | 78 | 74 |
| 12 | 75 | 69 |
| 13 | 77 | 74 |
| 14 | 75 | 70 |
| 15 | 77 | 77 |
| 16 | 76 | 72 |
| 17 | 70 | 73 |
| 18 | 70 | 73 |
| 19 | 77 | 74 |
| 20 | 77 | 73 |
| **Total** | **1,494** | **1,511** |
| **Mean** | **74.7** | **75.55** |

**c**

**Suppl. Table 2.** Instability

| **TCC#** | **7366** | **7365** |
|---|---|---|
| Your ID | Be Wo Cas9 | Be Wo dA11 |
| No. of meta. analy | 35 | 38 |
| No. of normal looking meta (1s) | 33 | 31 |
| No. of aberr. meta | 1 | 5 |
| No. of cells with breaks | 1 | 2 |
| No. of cells with fusions | 0 | 3 |
| Aneuploid/Tetra/Polyploid cells | 0 | 2 |
| No. of cells with Ganaphase morphology | 1 | 2 |

### Supplementary Figure 3

**a**

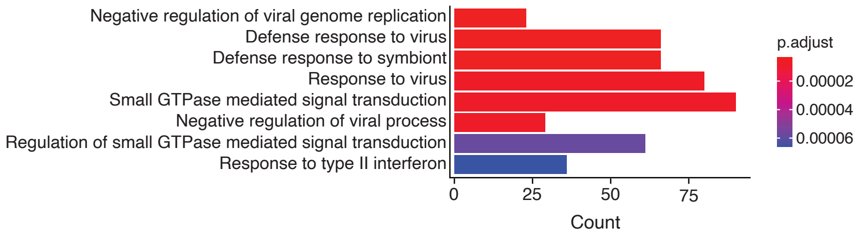


**b**

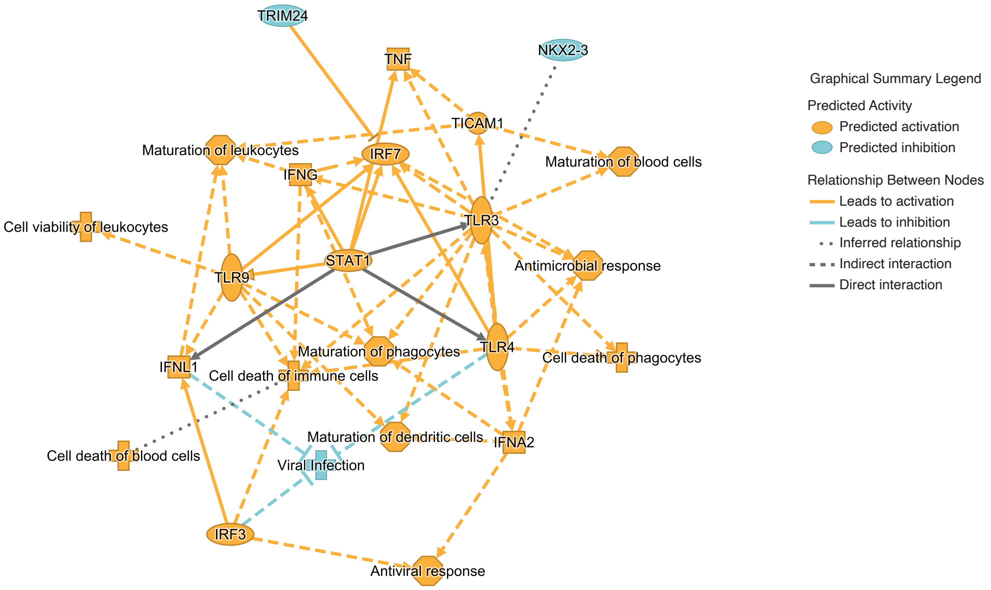

### Supplementary Figure 4

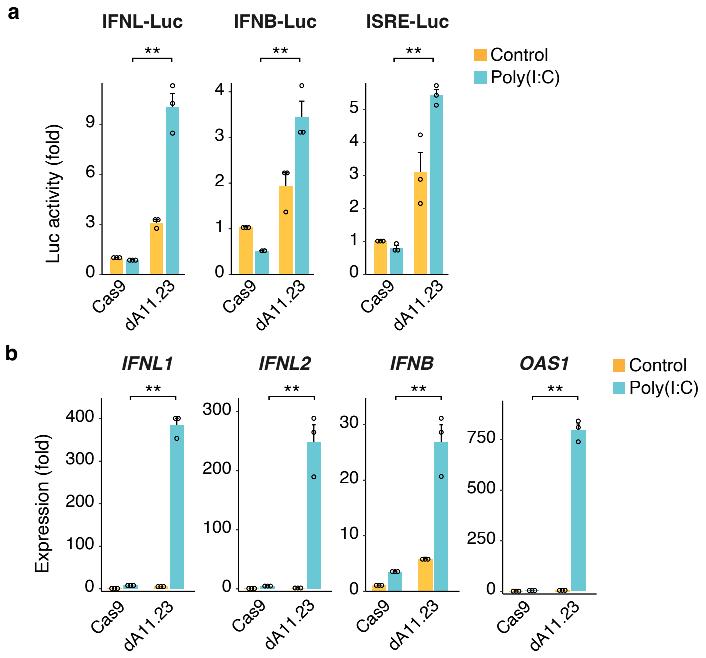

### Supplementary Figure 5

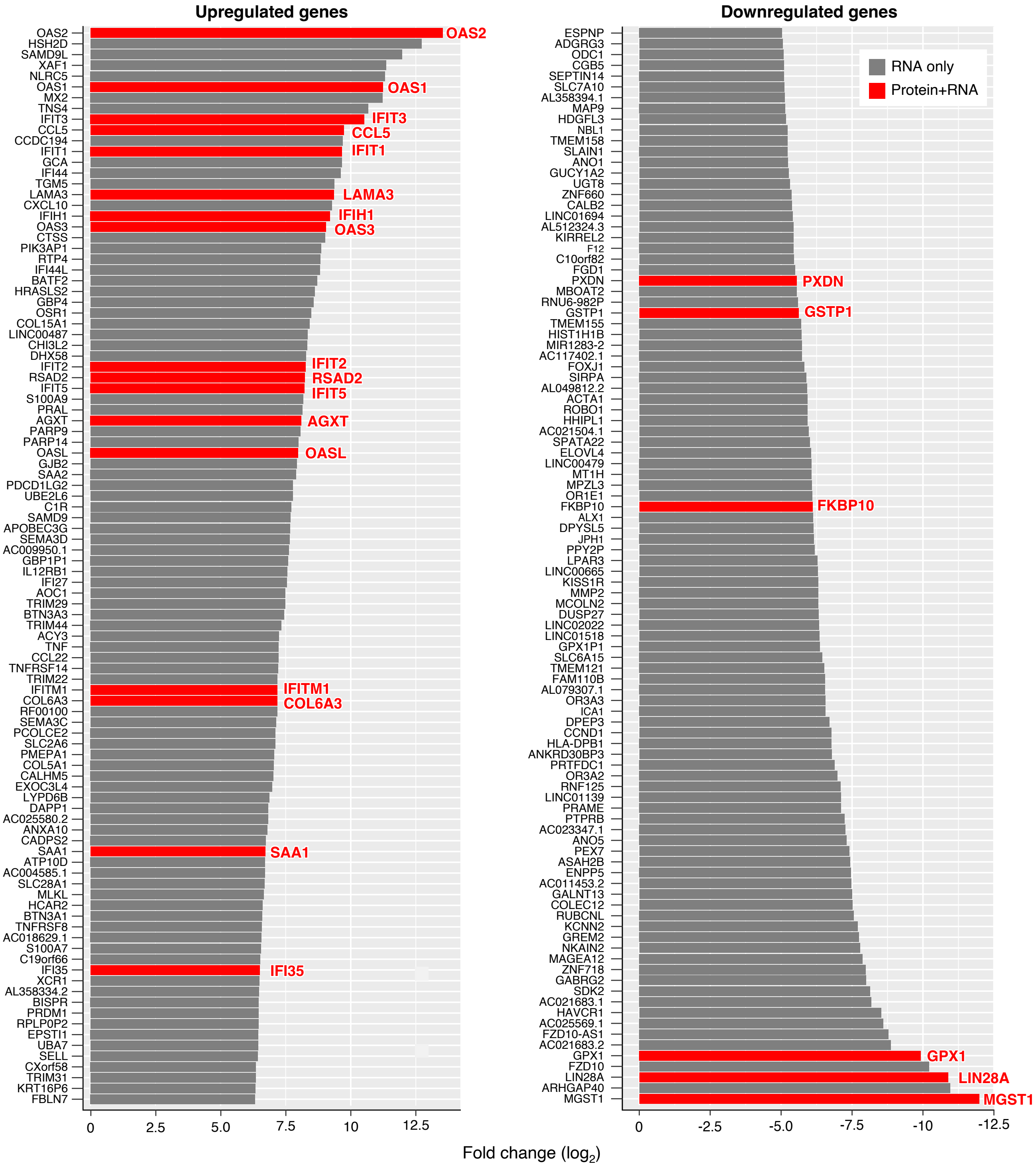

### Supplementary Figure 6

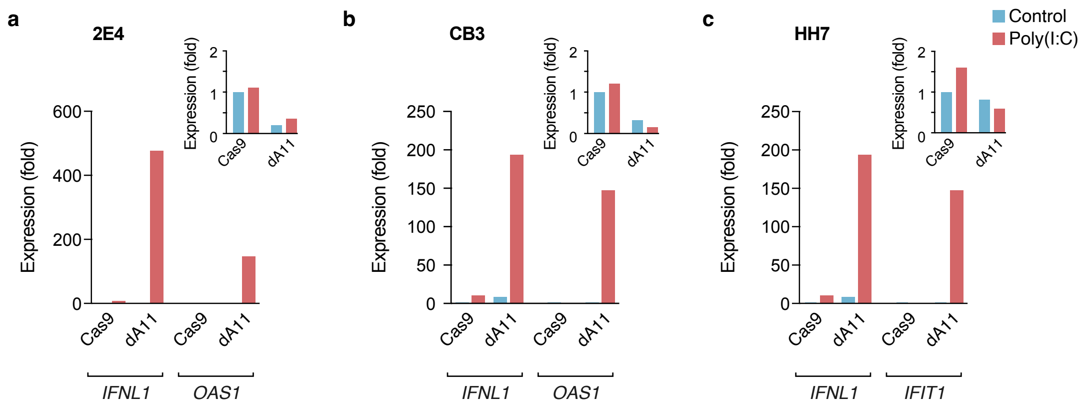

Supplementary Figure 7

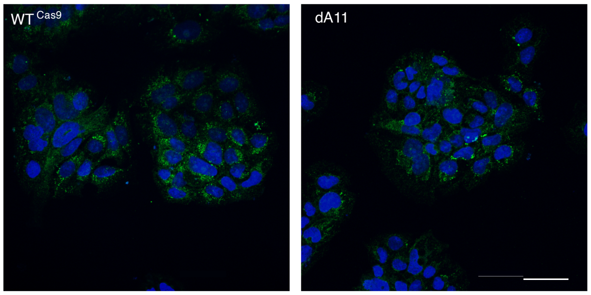

### Supplementary Figure 8

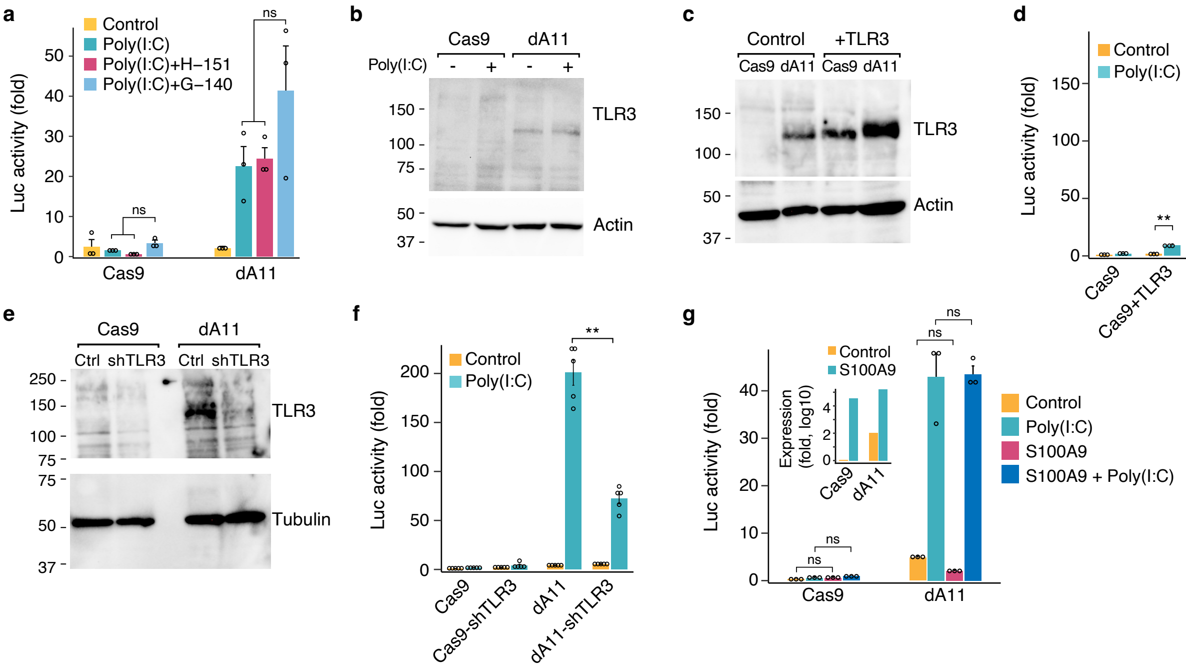

### Supplementary Figure 9

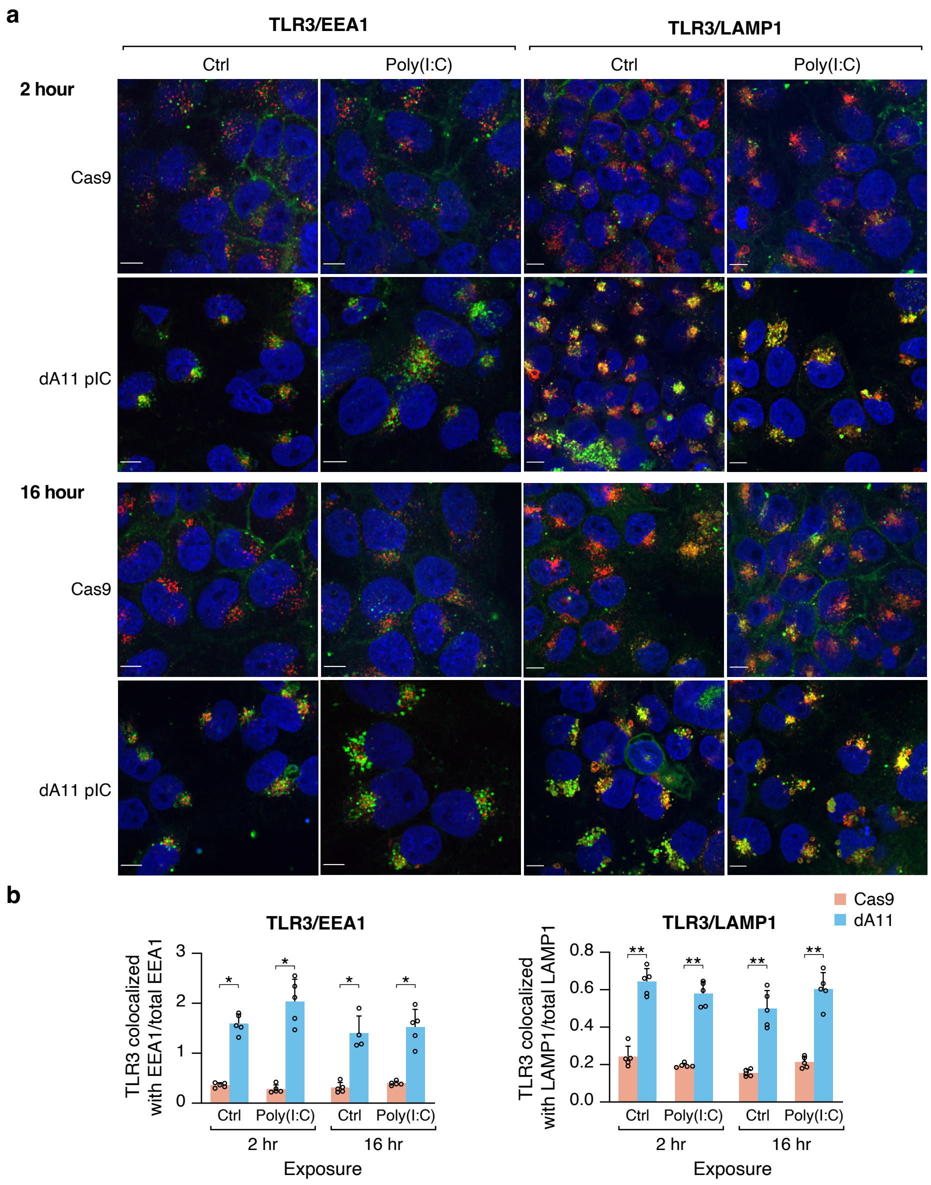

**Table S1**: Primers used in the study

| Primer | Sequence | Purpose |
|---|---|---|
| ATAC-1F | GATCGGTACCGGTTTTATGGCAGGATTTGG | cloning |
| ATAC-1R | GATCCTCGAGCCCTTCCTTCTAGTGCGTGA | cloning |
| ATAC-2F | GATCGGTACCGGGGAGCCAACACCTTATCT | cloning |
| ATAC-2R | GATCCTCGAGAGGCAAGTGGATCACCTGAG | cloning |
| ATAC-3/4F | GATCGGTACCACAGTGCCCGGCTAATTTTT | cloning |
| ATAC-3/4R | GATCCTCGAGCAGTGGCACAGTCTCAGCTC | cloning |
| ATAC-5F | GATCGGTACCAACTCCTGGCCACAAGTGAT | cloning |
| ATAC-5R | GATCCTCGAGTTATGCCCCTGCACTCTAGC | cloning |
| ATAC-6F | GATCGAGCTCCCCAGCCAAGATGTTTTTCT | cloning |
| ATAC-6R | GATCCTCGAGCTGCAACTTCTCTGCCTCCT | cloning |
| ATAC-7F | GATCGGTACCCAGTGGCTCACGCATGTAAT | cloning |
| ATAC-7R | GATCCTCGAGTGTGTGTTGGGACTGAAAGAA | cloning |
| ATAC-8F | GATCGGTACCGGAGGTTGGCAACACAAAAT | cloning |
| ATAC-8R | GATCCTCGAGGGATGGGCACAAAAACTAGG | cloning |
| ATAC-9F | GATCGCTAGCCATGGTGAGATCCCTGACTA | cloning |
| ATAC-9R | GATCCTCGAGGCACAATCTCGGCTCACTG | cloning |
| ATAC-10F | GATCGGTACCCCTGGAAGTATCGCCACCT | cloning |
| ATAC-10R | GATCCTCGAGTTGCAGTTCATCCACAAGGA | cloning |
| ATAC-11F | GATCGGTACCGCACCAGTTAAGAGGGATGC | cloning |
| ATAC-11R | GATCCTCGAGTTCCCAGAACCAGTCCAGTT | cloning |
| ATAC-11F_s | GATCGGTACCGCCTGTAATCCCAGCACTTT | cloning |
| ATAC-11R_s | GATCCTCGAGAGCGAGACTCCGTCTCAAAA | cloning |
| ATAC-12F | GATCGGTACCCCACATCGCTGGCTAATTTT | cloning |
| ATAC-12R | GATCCTCGAGTCACTCTGTCACCCAAGCTG | cloning |
| ATAC-12F_s | GATCGGTACCGCAGCATGTCACATCCAAAG | cloning |
| ATAC-12R_s | GATCCTCGAGTGGCAAGATTAGGGATGAGG | cloning |
| ATAC-13/14F | GATCGGTACCGGCATTGCATTGACTCTGAA | cloning |
| ATAC-13/14R | GATCCTCGAGACCCCGGGTATATCCAAAAG | cloning |
| ATAC-15F | GATCGGTACCCCAGCCTGTTTTCTGCATTT | cloning |
| ATAC-15R | GATCCTCGAGCTGCTCTCGAACTCCTGACC | cloning |
| ATAC-16F | GATCGGTACCTGTTTGTGTGTGCGACAATG | cloning |
| ATAC-16R | GATCAAGCTTCTAGATGGCCACAGCCTGA | cloning |
| ATAC-17F | GATCGAGCTCAGAGAGGCCAGTGTGGATGT | cloning |
| ATAC-17R | GATCCTCGAGTGGAAGGTTTTCATTGACGA | cloning |
| ATAC-17F_s | GATCGGTACCTGTGCCAAGCAAGATACAGG | cloning |
| ATAC-17R_s | GATCCTCGAGTGGTAGGAACCAACAAGGAAA | cloning |
| ATAC-18F | GATCGGTACCTGGATCAAACTGGTGCAACT | cloning |
| ATAC-18R | GATCCTCGAGCCCCCGTCTCTACCAAAAAT | cloning |
| ATAC-19F | GATCGGTACCCAGTTAATTTGCAGACCACCAA | cloning |
| ATAC-19R | GATCCTCGAGATTGGCTGTCGCCTGTAATC | cloning |
| IFNL1-F | GATCGAGCTCACCCCCAGAGGTTCTCATCT | cloning |
| INFL1-R | GATCGCTAGCGGCTAAATCGCAACTGCTTC | cloning |
| hOAS1-F | TGTCCAAGGTGGTAAAGGGTG | RT-qPCR |
| hOAS1-R | CCGGCGATTTAACTGATCCTG | RT-qPCR |
| hDDX60-F | CAGCTCCAATGAAATGGTGCC | RT-qPCR |
| hDDX60-R | CTCAGGGGTTTATGAGAATGCC | RT-qPCR |

| Primer | Sequence | Purpose |
|---|---|---|
| hOAS3-F | GAAGGAGTTCGTAGAGAAGGCG | RT-qPCR |
| hOAS3-R | CCCTTGACAGTTTTCAGCACC | RT-qPCR |
| hIFIH1-F | TCGAATGGGTATTCCACAGACG | RT-qPCR |
| hIFIH1-R | GTGGCGACTGTCCTCTGAA | RT-qPCR |
| hIFIT3-F | AAAAGCCCAACAACCCAGAAT | RT-qPCR |
| hIFIT3-R | CGTATTGGTTATCAGGACTCAGC | RT-qPCR |
| hIFIT3-F | AAAAGCCCAACAACCCAGAAT | RT-qPCR |
| hIFIT3-R | CGTATTGGTTATCAGGACTCAGC | RT-qPCR |
| hIFNL1-F3 | AATTGGGACCTGAGGCTTCT | RT-qPCR |
| hIFNL1-R3 | GTGAAGGGGCTGGTCTAGG | RT-qPCR |
| hIFNL2-F | AATTGTGTTGCCAGTGGGGA | RT-qPCR |
| hIFNL2-R | GCGACTGGGTGGCAATAAAT | RT-qPCR |
| hS100A9-F | GGTCATAGAACACATCATGGAGG | RT-qPCR |
| hS100A9-R | GGCCTGGCTTATGGTGGTG | RT-qPCR |
| hTLR3-F | GCCTTCTGCACGAATTTGAC | RT-qPCR |
| hTLR3-R | TCCAGCTGAACCTGAGTTCC | RT-qPCR |
| MCS-F | TCGAAGCTAGCTCTAGACTCGAGA | Cloning into pLV-hsp68 |
| MCF-R | CGATCTCTCGAGTCTAGAGCTAGCT | Cloning into pLV-hsp68 |
| VSV-F | TGCAAGGAAAGCATTGAACAA | RT-qPCR |
| VSV-R | GAGGAGTCACCTGGACAATCACT | RT-qPCR |
| GAPDH-F | GAAGGTCGGAGTCAACGGATTT | RT-qPCR |
| GAPDH-R | GAATTTGCCATGGGTGGAAT | RT-qPCR |
| YWHA-F | CTGAACTCCCCAGAGAAAGC | RT-qPCR |
| YWHA-R | CCGATGTCCACAATGTCAAG | RT-qPCR |
